# Preparing for the unknown: How working memory provides a link between perception and anticipated action

**DOI:** 10.1101/2022.01.25.477681

**Authors:** Marlene Rösner, Melinda Sabo, Laura-Isabelle Klatt, Edmund Wascher, Daniel Schneider

## Abstract

What mechanisms underlie the transfer of a working memory representation into a higher-level code for guiding future actions? Electrophysiological correlates of attentional selection and motor preparation processes within working memory were investigated in two retrospective cuing tasks. In the first experiment, participants stored the orientation and location of a grating. Subsequent feature cues (selective vs. neutral) indicated which feature would be the target for later report. The oscillatory response in the mu and beta frequency range with an estimated source in the sensorimotor cortex contralateral to the responding hand was used as correlate of motor preparation. Mu/beta suppression was stronger following the selective feature cues compared to the neutral cue, demonstrating that purely feature-based selection is sufficient to form a prospective motor plan. In the second experiment, another retrospective cue was included to study whether knowledge of the task at hand is necessary to initiate motor preparation. Following the feature cue, participants were cued to either compare the stored feature(s) to a probe stimulus (recognition task) or to adjust the memory probe to match the target feature (continuous report task). An analogous suppression of mu oscillations was observed following a selective feature cue, even ahead of task specification. Further, a subsequent selective task cue again elicited a mu/beta suppression, which was stronger after a continuous report task cue. This indicates that working memory is able to flexibly store different types of information in higher-level mental codes to provide optimal prerequisites for all required action possibilities.

**Highlights:**

• Selectively cueing features results in an overall performance benefit
• Feature-based attention is sufficient to form a prospective motor plan
• Prospective motor preparation can be initiated ahead of task specification
• Retro-active task specification leads to forming of higher-level action codes
• Different tasks requirements result in different prospective action plans

## 1. Introduction

In everyday life, we are frequently required to respond to stimuli that are present in our environment. However, what feels natural and automatic requires a complex cascade of cognitive processes, ranging from sensory perception via motor planning to motor execution. Working memory plays a central role in these cognitive processes, as it can be defined as a cognitive stage for the interface between perceptual information, higher-level cognitive operations, and goal-directed actions. Based on two working memory experiments and neural oscillations measured by means of the EEG, this study examined the role of prospective motor plans for the focusing of attention within working memory.

Traditionally, perception and action have been considered to occur in a capacity-limited, strictly serial fashion (Pashler, 1994). In this view, attention acts as a kind of spotlight by enhancing the mental representation of relevant information and potentially suppressing neural activity related to irrelevant content. Analogously, during the storage of visuo-spatial information in working memory, we can focus attention on certain stored content and thereby generate a prioritized representational state, protecting the attended information against decay and interference (Bays & Taylor, 2018; Makovski et al., 2008; Matsukura et al., 2007; Pertzov et al., 2013). However, this concept of an attentional bias on information stored in working memory fails to consider the goal-directedness of human information processing in every-day life. What we store in working memory is not only the mental representation of past sensory input, but also information about what current goal we are pursuing or what future action or mental operation we want to perform. Thus, working memory contents cannot only be seen as more or less precise copies of sensory information, but rather as mental representations that can be used to guide future action (for review: Nobre & Stokes, 2019; Olivers & Roelfsema, 2020).

In line with this account, earlier studies have shown that, just as storing sensory representations, working memory can also store higher-level representations of to-be-executed actions (e.g. Behmer & Fournier, 2014; Gallivan et al., 2016; Schneider et al., 2020; Zickerick et al., 2020). The attentional selection of certain working memory content could be the result of linking a sensory representation to such a representation of a to-be-executed action (Olivers & Roelfsema, 2020). In this regard, the measurement of neural oscillations in the EEG as a correlate for motor planning can provide valuable information. It was shown that oscillatory power in the mu (∼10-14 Hz) and beta frequencies (∼15-30 Hz) can track the preparation of a motor response (Leocani et al., 1997; Pfurtscheller et al., 2000; Zhuang et al., 1997). For example, when participants can acquire explicit knowledge about a certain action sequence across an experiment, this knowledge about which action to-be-executed next is reflected in a suppression of mu oscillatory power over motor areas contralateral to the side of response (Zhuang et al., 1997). In a comparable way, an investigation by Schneider, Barth, & Wascher (2017) made use of oscillatory effects in mu/beta power as correlates of the creation of motor representations during the storage interval of a visual working memory task.

Participants had to store three different visual items and were then cued to focus on one, two or three items for later report by means of a right-handed movement of the computer mouse. Following these cues, the authors observed a suppression of mu and beta oscillatory power with an estimated source in the pre-motor and motor cortex contralateral to the side of the to-be-executed action. Just like in the motor learning experiment by Zhuang and colleagues (1997), this effect was only found when participants gained explicit knowledge about the action to-be-executed next (i.e., when only one item was cued) and it appeared clearly prior to the presentation of the memory probe demanding the actual response. In a comparable way, van Ede and colleagues (2019) used non-spatial (color) information to cue relevant working memory content. They orthogonally manipulated the lateralized position of the relevant visual item in a memory array (left vs. right target) and the required motor response (button press with left vs. right index finger). Hemispheric asymmetries in alpha oscillations related to attentional selection (Foxe & Snyder, 2011; Händel et al., 2011; Myers et al., 2015; Sauseng et al., 2005; Schneider et al., 2016) and in mu and beta oscillations related to planning a left- vs. right-handed response could thus be fully separated from one another. It was shown that these types of oscillatory effects were related to spatially distinct areas of the brain but occurred in the same time frame. These results highlight that working memory can store sensory information along with corresponding motor plans and that both kinds of memory codes can be selected in support of future behaviour (see also: Boettcher et al., 2021).

So far, the mentioned investigations suggested that correlates of motor planning like the suppression of mu/beta oscillatory power result from selecting an individual object stored in working memory for action control. However, in order to establish a general role of motor planning processes during the selection and storage of working memory content, it is necessary to investigate whether these principles also apply to the selection of individual features of stored objects. Therefore, as a first step, the current study investigated to what extent feature selection in working memory also involves the selection and storage of prospective motor plans. For this purpose, participants had to store both the location and orientation of a single object in working memory and were then cued to focus on one of these features for later report. We expected that both selective cues should lead to a more accurate report relative to a neutral cue condition without an attentional bias towards either feature.

Furthermore, if selecting a visual feature from an object stored in working memory also involved motor planning (or the selection of an associated motor code), we should observe stronger suppression of oscillatory mu and beta power with an estimated source in the pre-motor cortex contralateral (but not ipsilateral) to the to-be-executed response for the selective cue conditions relative to the neutral condition. This effect should occur together with a stronger suppression of alpha power over posterior visual areas following the selective cues that was already observed in the context of the retrospective selection of visual object features (see Hajonides et al., 2020; Niklaus et al., 2017; Sasin & Fougnie, 2020; Ye et al., 2016).

In a second experiment, we wanted to build on this by asking what conditions must be met for a prospective motor code to get created in working memory. For example, it is possible that a motor representation can be created only if it is clearly defined that the next operation in a working memory task requires dealing with the stored information based on a certain movement (e.g., adjusting a memory probe orientation or moving the hand or gaze to a specific position). So far, this was always the case in the studies that used mu and beta oscillations as a correlate of motor planning during working memory storage (Boettcher et al., 2021; Schneider et al., 2017; van Ede et al., 2019). Such clear knowledge about the task to be executed next has been shown to improve working memory accuracy (Printzlau et al., 2019) and modulate the way the storage of visual working memory content is represented in the EEG signal (Fahrenfort et al., 2017). However, it is also possible that a motor representation of task-relevant working memory content is created even if the upcoming task has not yet been clearly defined. Working memory could then flexibly select from the stored mental representations of a given content the one(s) best suited to support each subsequent operation. In the present study, participants therefore had to select the relevant information in working memory on the basis of a feature cue (location vs. orientation vs. neutral) and later use it either for a continuous report task (adjusting a memory probe to the target feature by moving the computer mouse) or a recognition task (match vs. mismatch decision based on a memory probe). Only a second retro-cue (continuous report vs. recognition) or the later memory probe display (following a neutral task cue) indicated which of the two tasks should be performed based on the selected information. We hypothesized that a motor code of the selected information should be pre-emptively created in working memory even if the task to be performed has not yet been clearly defined. In this case, a greater suppression of mu and beta power in contralateral sensorimotor and pre-motor cortex after the selective feature cues (i.e., before the definition of the task to be performed) relative to the neutral condition should be evident as a correlate of motor selection. A further selection of the mental representations useful for the to-be-executed task should then take place after the (second) task retro-cue.

Compared to the recognition task which should first require comparing the memory probe to the relevant visual feature before selecting a certain response, cuing the continuous report task should entail a motor-planning process ahead of memory probe presentation, reflected by a stronger suppression of mu and beta power in contralateral sensorimotor and pre-motor cortex. These results would argue for a flexible use of sensory and motor codes in working memory according to the requirements of the tasks to be performed.

## 2. Materials and methods

### 2.1. Participants

Twenty-four participants took part in the first experiment (15 females), who were between 20 and 30 years old (*M* = 24.13, *SD* = 3.03) and right-handed (as assessed with the Edinburgh Handedness Inventory: Oldfield, 1971). Twenty-six participants took part in the second experiment. Due to performance below chance level (n = 3) and one participant actually being left-handed, four participants had to be excluded. The final sample in the second experiment thus consisted of 22 right-handed participants (age: *M* = 23.09, *SD* = 2.43, range = 19-27, 19 female). None of the participants from the first and second experiment reported suffering from any neurological or psychological disorder. Participation was compensated with 10 Euros/hour or with course credits (for psychology students). The experiments were approved by the local ethics committee of the Leibniz Research Centre for Working Environment and Human Factors (Dortmund, Germany) and were conducted in accordance with the Declaration of Helsinki.

### 2.2. Stimuli and procedure

The experiments took place in an electrically shielded, dimly lit chamber. Participants were seated with a viewing distance of 150 cm from the 22-inch CRT monitor (1024x768 pixels), which had a 100 Hz refresh rate. A ViSaGe MKII Stimulus Generator (Cambridge Research Systems, Rochester, UK) controlled stimulus presentation.

Prior to the actual experiments, participants performed a short training and three practice blocks to get acquainted with the tasks. During the training, a grey circle (luminance = 20cd/m^2^, RGB = [R54 G54 B54], diameter = 4.4°) was presented on the left side of the screen. A randomly oriented Gabor grating (size = 1.3°, spatial frequency = 4.25 cycles per degree, phase = 180°, deviation = 0.25°) was shown on the perimeter of the circle on a random position. On the right side, a similar second circle with either one bar (indicating a position report) or two opposing bars (indicating an orientation report) was shown. The participants’ task was to adjust the second circle, using lateral mouse movements, so that the bar(s) matched the position or the orientation of the Gabor grating. In cases when the adjustment took longer than 4 s, participants were instructed to respond faster in order to familiarize themselves with the time constrains of the later experiment. The training was complete when the average response error was below 18° for orientation and below 36° for location in 50 consecutive trials. Afterwards, participants performed three practice blocks (30 trials each), one for each condition of the experiment (see below).

#### 2.2.1. Experiment 1

Participants were instructed to memorize the orientation and location of an oriented Gabor grating (similar to the one in the training; 16 different possible locations 22.5° apart from each other; eight different possible orientations 22.5° apart from each other), depicted on the perimeter of an imaginary centrally located circle (diameter = 4.4°). After a 1500 ms delay interval, a retrospective feature cue indicated whether participants would have to recall the location (location cue: “P” for “Position”; one third of the trials) or orientation (orientation cue:”, “O”; one third of the trials) of the initially presented grating (see figure 1A). In one third of the trials, the feature cue contained no information about the to-be-reported feature (neutral cue, “X”). After another delay period, a highly salient distractor appeared in the centre of the screen followed by another delay interval. The visual distractor was included for facilitating decoding of the cued working memory representations (see Wolff et al., 2017) and therefore is not relevant for the current investigation. Finally, a memory probe, which consisted of a circle with either one marker (location report) or two opposing markers (orientation report) at a randomly chosen position appeared. Upon memory probe presentation, participants had to adjust the marker(s) position to indicate the cued feature (i.e., the grating’s location or orientation). Following a neutral feature cue, the type of memory probe (one mark vs. two marks) indicated the target feature. Once the probe was correctly adjusted, each answer had to be confirmed by a button press within 4 s. Otherwise the trial was considered incomplete. The task consisted of 720 trials (240 per feature cue condition) and the whole experiment (including the EEG cap preparation) lasted around 3.5 hours.

**Figure 1.**
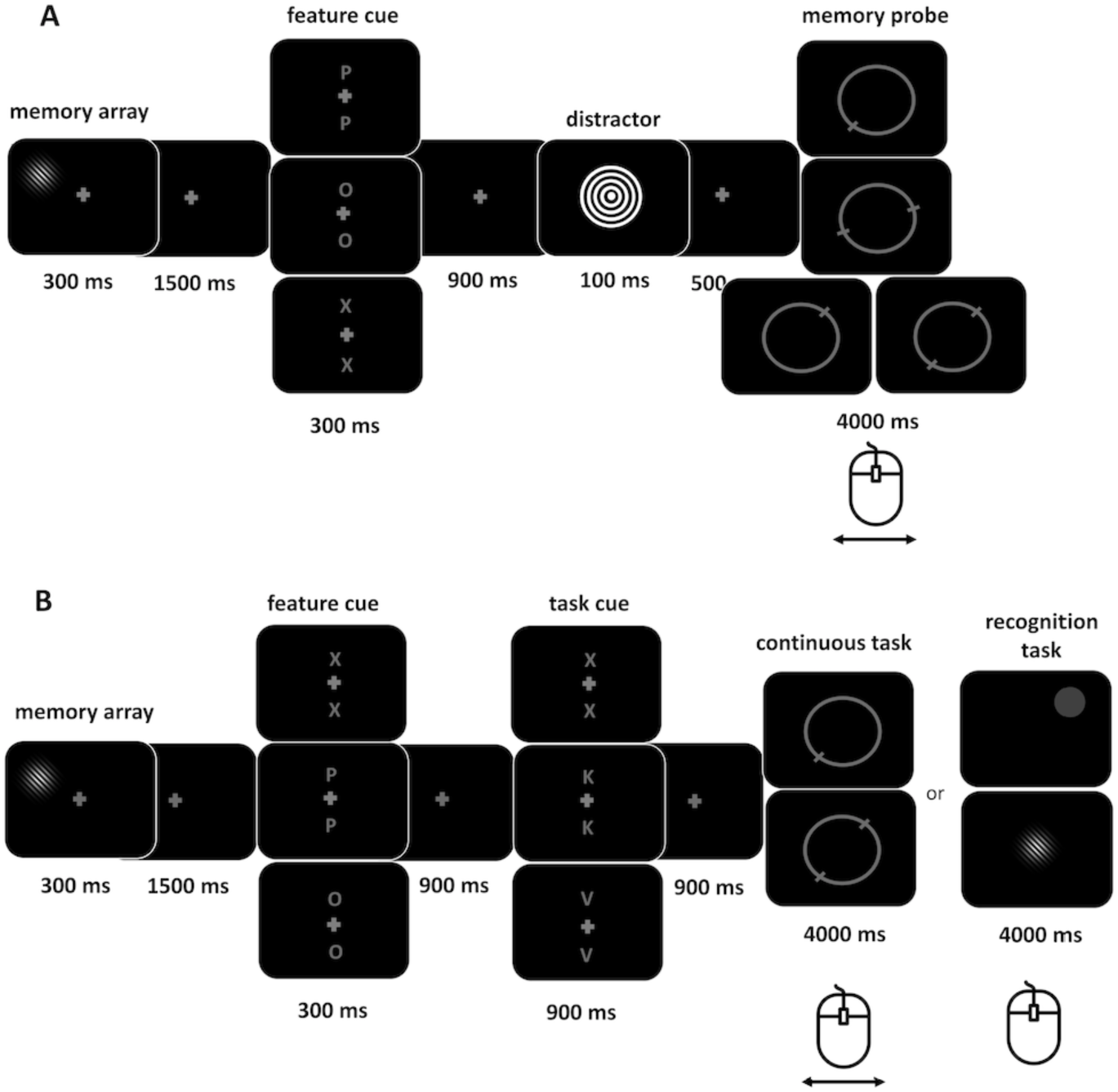
Experimental design. Panel A depicts the procedure of Experiment 1. Each trial began with the presentation of an oriented Gabor grating whose location and orientation had to be memorized. After a delay interval, a selective feature cue (P = position, O = orientation, X = neutral) indicated whether participants had to reproduce the orientation or location. Finally, after presentation of a visual distractor stimulus, a memory probe had to be adjusted so that it matched the target feature. The design of Experiment 2 is depicted in panel B. Here, the feature cue was followed by a task cue (X = neutral, K = continuous report task, V = recognition task) indicating whether a continuous report task (as in Experiment 1) or a recognition task had to be performed. In the recognition task, either a probe location or a probe orientation was presented, and participants indicated by mouse click whether it matched the target information from the memory array.

**Figure 2.**
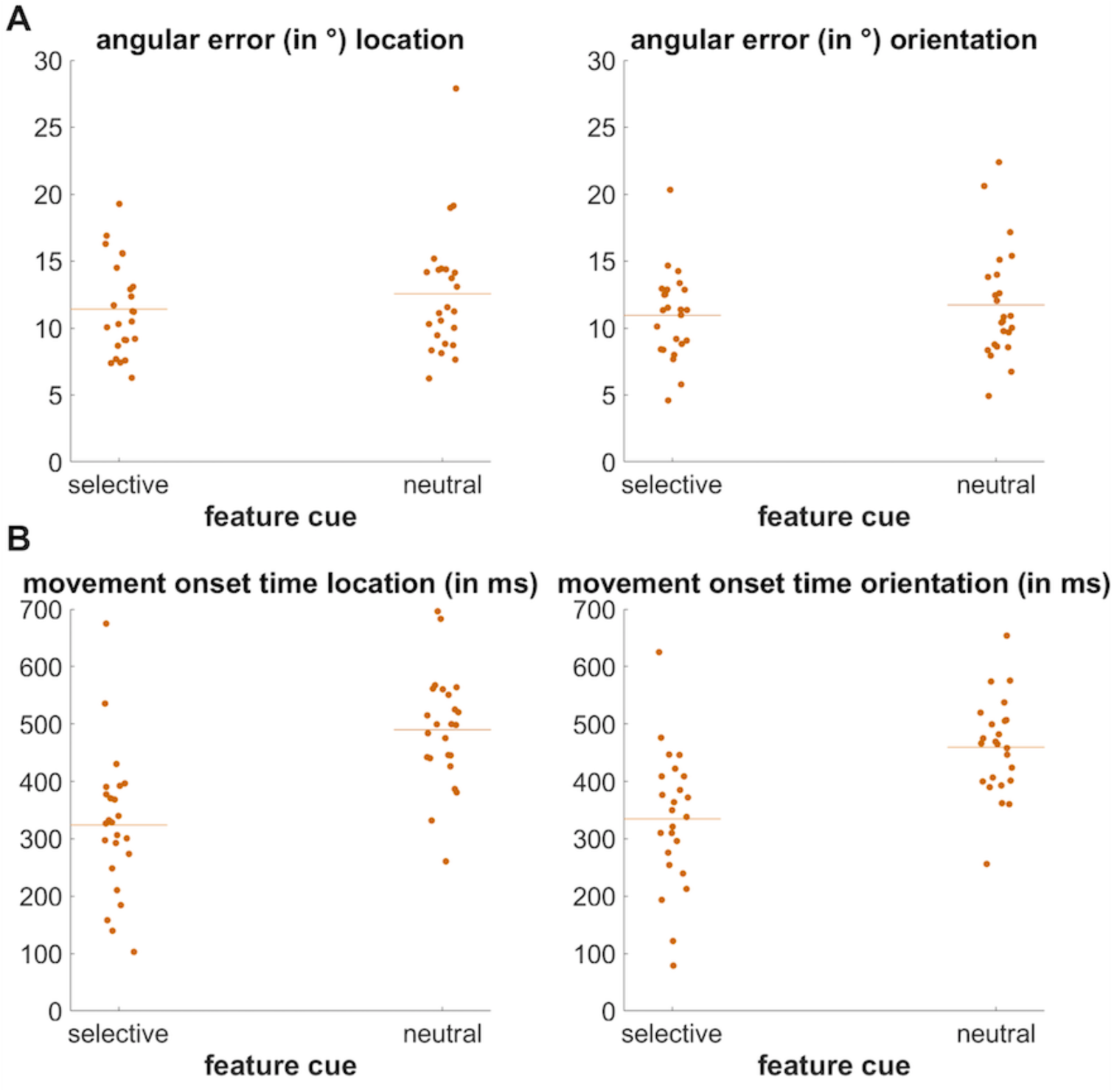
Behavioural results of Experiment 1. Panel A shows the angular error for location feature cues vs. neutral cue conditions (left) and for the orientation feature cue vs. neutral cue conditions (right). The coloured dots depict the mean angular error of each participant and the horizontal line depicts the mean across participants. Panel B represents comparisons of the time to mouse movement onset between the above-mentioned conditions.

#### 2.2.2. Experiment 2

The second experiment differed from the first one in the sense that participants had to perform one of two possible tasks: either the same location/orientation continuous report task as in Experiment 1 (50% of the trials) or a recognition task (50% of the trials). After memorizing the location and orientation of the grating presented in the memory array, a retrospective feature cue (location cue, orientation cue or neutral feature cue) indicated which feature was going to be relevant at the end of the trial. The delay interval after the feature cue was followed by a second retro-cue indicating whether participants had to adjust the location or orientation of the probe or to compare a presented feature with the cued feature from the memory array. When the continuous report task was cued (one third of the trials; “K”, K = kontinuierlich; i.e., the German word for “continuous”; see figure 1B) the memory probes were identical to the first experiment. When the recognition task was cued (one third of the trials; “V”, V = Vergleich; i.e., German for “comparison”) the probe displayed just one feature. The probe location was indicated by a filled grey circle presented at a given location on an imaginary circle. Participants had to indicate by pressing the left or right button on the computer mouse (right hand) whether the position of the grey circle matched the initial position of the memory item. In orientation-recognition trials, a grating with a specific orientation was presented in the centre of the screen. Again, participants had to indicate with the computer mouse buttons whether this was the same or different orientation as presented in the memory array. The probe matched the memory array in half of the recognition task trials. The assignment of responses to the computer mouse buttons was counterbalanced across participants. In one third of the trials, the task cue was neutral, and participants could infer which task they should perform by looking at the probe display. When the probe was a circle with one or two marks on it, they had to perform the continuous report task. When it was either a filled grey circle or a centrally presented grating, the recognition task had to be performed. The combination of feature and task cues resulted in nine conditions (3 x 3 experimental design) with 80 trials each. The experiment consisted of 720 trials in total and took about 4 hours, including the preparation of the EEG cap.

### 2.3. Behavioural Analysis

#### 2.3.1 Experiment 1

Only complete trials (including a button press following location/orientation report) were included in the analyses and two parameters were considered: the angular error (calculated as the difference between the original orientation/location of the grating and the reported value) and the time to mouse movement (i.e., the time required for starting response initiation). For analysis of the angular error, conditions were compared against the respective neutral retro-cue condition with a *t*-test separately for each feature. More specifically, trials with a location feature cue were compared to neutral trials with a location probe and orientation feature cue trials were compared to neutral trials with a later orientation probe. Since location and orientation adjustment featured different maximal angular error values (location: 180°; orientation: 90°), angular error analysis was done separately for each feature. Time to mouse movement onset was analysed with a repeated measures analysis of variance (rm-ANOVA) with the factors *feature* (location vs. orientation) and *cue type* (selective vs. neutral).

#### 2.3.2. Experiment 2

Parameters for the analysis of working memory accuracy and response initiation in the continuous report task were the same as in Experiment 1: the angular error and time to mouse movement onset. Only complete trials were included in all analyses. For the recognition task, the percentage of correct responses and response times were analysed. Here, responses were considered from 150 - 4000 ms after memory probe presentation. Trials with responses prior to this interval were not included in the analyses (premature responses). Responses were considered as erroneous when there was a wrong button press or no button press within 150 - 4000 ms (misses). For analysis of the angular error, separate rm-ANOVA were run with the factors *feature cue* (selective vs. neutral) and *task cue* (selective vs. neutral). All other parameters (continuous task: time to mouse movement onset; recognition task: percentage of correct responses, response times) were analysed with a rm-ANOVA with the factors *feature* (location vs. orientation), *feature cue* (selective vs. neutral) and *task cue* (selective vs. neutral).

### 2.4. EEG recording and pre-processing

EEG data were recorded using a 128 Ag/AgCl passive electrode cap (Easycap GmbH, Herrsching, Germany) with a 10/20 configuration (Pivik et al., 1993). Data were recorded with a sampling rate of 1000 Hz and amplified by a NeurOne Tesla AC-amplifier (Bittium Biosignals Ltd, Kuopio, Finland). During data acquisition, an online 250 Hz low-pass filter was applied, which was chosen to be well below the Nyqist frequency in order to prevent aliasing. Impedances were kept below 20 kΩ. The FCz electrode was chosen as reference and the AFz as ground electrode.

EEG data were analysed using MATLAB (R2021a) and the EEGLAB toolbox (v.14.1.2, Delorme & Makeig, 2004). As the first pre-processing step, data were 1 Hz high-pass (roll-off= 48dB/oct, half-amplitude cut-off: 1 Hz, half-power cut-off: 1.1 Hz) and 40 Hz low-pass filtered (roll-off = 48dB/oct, half-amplitude cut-off: 40 Hz, half-power cut-off: 36.4), using an infinite impulse response (IIR) Butterworth filter (*pop_basicfilter* function included in the ERPLAB toolbox; Lopez-Calderon & Luck, 2014). Prior research has shown that 1-2 Hz high-pass filtering of the continuous data can optimize ICA decomposition (see Winkler et al., 2015). The 40 Hz low-pass filter was applied to exclude high-frequency fluctuations in the EEG signal that were not of interest for the current investigation (> 30 Hz; e.g., line noise).

After filtering, the data were down sampled to 250 Hz to decrease further processing time. Channels containing a high level of artifacts were excluded using an automated channel rejection procedure (*pop_rejchan* function included in EEGLAB, probability threshold = 5 SD; kurtosis threshold = 10). On average, 5.79 channels were excluded per participant (*SD* = 3.75, range = 0-15) in Experiment 1. In Experiment 2, 6.92 channels per participant were excluded on average (*SD* = 4.6, range = 1-15).

Next, data were re-referenced to a common average and were then separated into epochs time-locked to the memory array (Experiment 1: 1000 ms before and 6100 ms after the memory array; Experiment 2: 1000 ms before and 7400 ms after the memory array). Trials containing extreme fluctuations were excluded with an automated artifact rejection procedure (*pop_autorej* function included in EEGLAB; threshold = 500 µV, probability threshold = 5 SD, max. % of rejected trials per iteration: 5%). On average, 664.42 trials remained for the ICA (*SD* = 36.86, range = 592 - 711) in Experiment 1 and 672.84 trials (*SD* = 34.06, range = 592 - 715) in Experiment 2.

ICA was conducted on the rank-reduced data (number of channels minus 1, i.e., to account for the rank deficiency introduced by re-referencing to average reference) and components reflecting artifacts (horizontal and vertical eye movements, blinks, generic data discontinuities) were identified using ADJUST (Mognon et al., 2011). In addition, single-equivalent current dipoles were fitted on the ICs based on a spherical head model using the dipfit-plugin of the EEGLAB toolbox. ICs with a residual variance exceeding 50% regarding their dipole solution and those ICs identified by ADJUST were excluded. Overall, 50.33 components (*SD* = 11.30, range: 32-73) were excluded in Experiment 1. In Experiment 2, 51.05 components (*SD* = 11.39, range: 32-77) were excluded.

Trials still containing extreme fluctuations were identified and excluded through a second automated artifact rejection procedure (threshold = 1000 µV, probability = 5 SD, max. % of trials rejected per iteration = 5%). On average, 195.13 trials per experimental condition (*SD* = 16.46, range = 166 – 228.67) (Experiment 1) remained. In Experiment 2, 67.48 (*SD* = 5.6, range = 58.33 – 76.11) trials per experimental condition remained on average. Finally, rejected channels were interpolated based on a spherical spline algorithm (*eeg_interp* function included in EEGLAB).

### 2.5. Channel-based analysis

Spectral power was computed by convolving complex Morlet wavelets with each trial of the EEG data. Frequencies between 4 and 30 Hz were included, which increased in logarithmic steps of 52. The width of the Gaussian, defined by the number of cycles, increased linearly by a factor of 0.5, resulting in three cycles at the lowest and 11.5 cycles at the highest frequency. An interval of 200 ms before memory array onset (-200 to 0) was used as spectral baseline. The resulting epochs contained 200 time points ranging from -1000 ms before to 6000 ms after memory array onset (Experiment 2: -1000 ms to 7400 ms after memory array onset). For both experiments, ERSPs were computed separately for each experimental condition (Experiment 1: feature cue conditions; Experiment 2: feature and task cue combinations). Depending on the particular research questions, averages were taken across conditions for further analysis.

Similar to Schneider et al. (2017), four clusters of electrodes were chosen to investigate to what extent the retrospective cuing of object features led to a modulation of oscillatory power prior to memory probe presentation: two clusters over the left (PO3, PO7, PPO5h, P7, P5) and right (PO4, PO8, PPO6h, P8, P6) parieto-occipital cortex and two clusters over the left (CP3, CCP5h, CCP3h, C3) and right (CP4, CCP6h, CCP4h, C4) sensorimotor cortex. As a first step, data were averaged across these clusters and across the two selective feature cue conditions. A cluster-based permutation procedure was performed in order to find the time window and frequency range in which selective and neutral feature cue conditions differed from each other. Since differences in this regard should appear following the retro-cue (the first retro-cue for Experiment 2), only time points between the retro-cue and the probe were included in the permutation procedure. Condition labels (selective vs. neutral feature cue) were randomly assigned to each dataset. This was repeated 1000 times. A two-sided within subject *t*-test was performed for each time-frequency data point on each iteration resulting in a time points (67) x frequencies (52) x permutations (1000) matrix (Experiment 2: time points (63) x frequencies (52) x permutations (1000)). For each permutation, the size of the largest time-frequency cluster with *p* < .05 was assessed. Differences between selective and neutral feature cues in the original data were considered significant, if the size of time-frequency cluster was larger than the 95^th^ percentile of the distribution of cluster sizes created by the permutation procedure.

Subsequently, a rm-ANOVA with the factors *condition* (location feature cue vs. orientation feature cue vs. neutral feature cue), *hemisphere* (left vs. right electrode clusters) and *caudality* (posterior vs. central electrode clusters) was performed on the identified time-frequency clusters. The three-way interaction of these factors was analysed to test whether the stronger contralateral (left-hemispheric) vs. ipsilateral (right-hemispheric) suppression in mu/beta oscillatory power we expected for the selective feature cue conditions was indeed stronger over the motor cortex (central electrode clusters) and not rather related to retro-cue effects on more posterior oscillatory patterns in the alpha and beta frequency ranges.

### 2.6. Independent component clustering

A clustering procedure on IC level, as implemented in the EEGLAB toolbox, was used to further isolate the mu/beta activity over the sensorimotor cortex from posterior alpha activity and to prove the sensorimotor source of the mu/beta suppression.

All parameters used for the clustering procedure were based on the approach by Schneider et al. (2017). However, the number of clusters resulting from the k-means clustering algorithm was changed to 24, due to the higher number of channels (resulting in a higher number of ICs). We chose this higher number of resulting clusters to guarantee a sufficient solution with many datasets contributing ICs to the mu/beta clusters, while still ensuring a good isolation of the mu/beta IC clusters from those reflecting more posterior alpha/beta oscillations. Only ICs with less than 20% residual variance regarding their dipole solution were included in the clustering procedure (see Schneider et al., 2017). Event-related spectral perturbations (ERSPs) for the individual ICs were calculated with the same parameters as for the channel-based analysis. Frequency spectra were computed using an FFT (fast-Fourier transform) algorithm. The clustering was based on estimated dipole locations (three dimensions), scalp distributions (10 dimensions), ERSPs between 4 and 30 Hz (10 dimensions) and spectral power between 4 and 30 Hz (10 dimensions). The number of dimensions define to what extent the different features contribute to the generation of the clusters (Onton & Makeig, 2006). As the IC dipole solution can only contribute three dimensions (x, y and z values), its relative contribution to clustering was weighted by a factor of 10 (see the *std_preclust* function included in EEGLAB). A k-means clustering algorithm separated 870 ICs into 24 clusters (Experiment 2: 835 ICs into 24 clusters). Furthermore, an individual IC was considered as an outlier when it was more than 3 SD away from any of the 24 cluster centroids (referring to the distance between the IC and the locations representing the centre of each IC cluster in the multidimensional feature space; see *pop_clust* function included in EEGLAB).

To further illustrate the estimated neural sources of the IC clusters used for further analyses (see Results section), we specified the MNI coordinates of the dipole centroid for each cluster. MNI coordinates resulted from the dipole fitting procedure where the channel locations were aligned with a spherical head model and an average MRI image from the Montreal Neurological Institute (MNI) database (average of 152 T1-weighted stereotaxic volumes; International Consortium for Brain Mapping/ICBM). Thus, each individual component was assigned a dipole with three coordinates (x,y,z), which can be mapped to a specific brain region via the average MNI brain template. Based on all ICs within a cluster, we then expanded this centroid point to a sphere with a radius that had the length of 1 *SD* referred to each of the three dipole coordinates. The statistical sources were defined by the number of grid points within this extended spatial sphere that belonged to a specific anatomical structure, divided by the number of all grid points (see figures 5, 7 and 8). This procedure was based on the *std_dipoleDensity* function implemented in the EEGLAB toolbox. For statistical analyses, ICs from subjects contributing several ICs to one of the clusters were averaged.

For Experiment 1, ERSP responses generated by the statistical sources from 19 participants (left-hemispheric cluster) and 20 participants (right-hemispheric cluster) were compared between the two selective feature cue conditions (location vs. orientation) and the neutral condition by a cluster-based permutation approach comparable to the one described for the channel-based analysis. For Experiment 2 (left-hemispheric cluster: 22 participants; right-hemispheric cluster: 23 participants), the same cluster-based permutation approach was used for analysing condition effects based on the first feature retro-cue (location vs. orientation vs. neutral) and the second task cue (continuous report vs. comparison vs. neutral).

### 2.7. Inferential statistics and effect sizes

When indicated by Mauchly’s test for sphericity, Greenhous-Geisser correction was applied (indicated by *ε*). Effect sizes for ANOVAs are indicated by partial eta squared (*η^2^_p_*) and by Cohens *d_z_* for within-subject *t*-tests. To prevent *p*-value inflation due to multiple comparisons, the false discovery rate (FDR) procedure (adjusted *p* values / *p_adj_* are reported in this regard; Benjamini & Hochberg, 1995) was used for post-hoc comparisons and to adjust for hidden multiplicity within rm-ANOVAs (the critical *p*-value or *p_crit_* is reported in this regard; Cramer et al., 2016). Additionally, the boundaries of the 95^th^ confidence interval (CI) surrounding the mean condition differences are reported.

## 3. Results

### 3.1. Behavioural results

#### 3.1.1. Experiment 1

Precision of working memory performance was assessed by the difference between the target feature value and the adjusted probe. This angular error (see figure 2A) for location adjustment was lower following a location feature cue than following a neutral feature cue, *t*(23) = -2.47, *p_adj_* = .043, *d_z_* = -0.50, CI 95% [-2.12 -0.19]. However, this effect was increased by on outlier in the neutral feature cue condition. After outlier exclusion (mean +/- 3 standard deviations), this effect could no longer be considered as statistically significant (*t(23) = -2.36, p_adj_* = .055, d*_z_* = -0.49, CI 95% [-1.55 0.10]). Similarly, the angular error for orientation adjustment trials did not differ between selective and neutral feature cues, *t*(23) = -1.88, *p_adj_* = .072, *d_z_* = -0.38, CI 95% [-1.64 0.08]. Speed of response initiation (indicated by time to mouse movement onset; see figure 2B) was accelerated by the selective feature cue, *F*(1,23) = 151.38, *p* < .001, *p_crit_* = .017, *η^2^_p_* = 0.87. Speed of location adjustment did not differ from orientation adjustment, *F*(1,23) = 1.74, *p* = .200, *p_crit_* = .05, *η^2^_p_* = 0.07. There was also an interaction of task and feature cue, *F*(1,23) = 11.85, *p* = .002, *p_crit_* = .033, *η^2^_p_* = 0.34.

Comparison of effect sizes revealed that time to response initiation was slightly more decreased after a selective compared to a neutral cue in location, *t*(23) = -11.48, *p_adj_* <.001, *d_z_* = -2.34, CI 95% [-195.83 -136.01], than in orientation adjustment trials, *t*(23) = -10.49, *p_adj_* < .001, *d_z_* = -2.14, CI 95% [-149.37 -100.16].

### 3.1.2. Experiment 2

For Experiment 2, precision of working memory performance (see figure 3A) in the continuous task was reliably increased for location adjustment by a selective feature cue, as revealed by a main effect of feature cue, *F*(1,21) = 14.26, *p* = .001, *p_crit_* = .017, *η^2^_p_* = 0.40, while there was no influence of the task cue on probe adjustment, *F*(1,21) = 0.37, *p* = .548, *p_crit_* = .05, *η^2^_p_* = 0.02. Orientation adjustment was not affected by any of the cues (feature cue: *F*(1,21) = 0.04, *p* = .845, *p_crit_* = .033, *η^2^_p_* < .01; task cue: *F*(1,21) < 0.01, *p* = .962, *p_crit_* = .05, *η^2^_p_* < 0.01).

**Figure 3.**
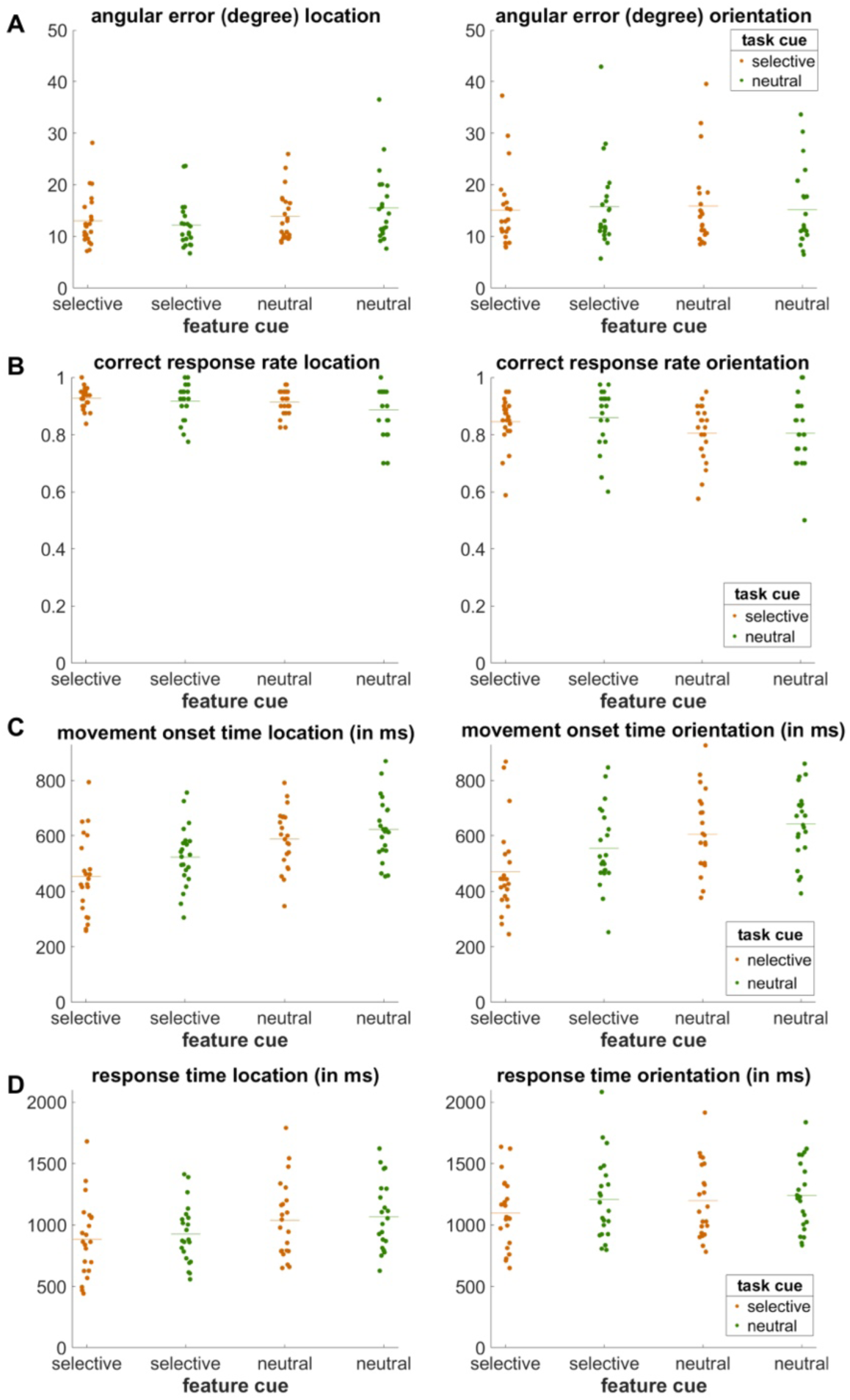
Behavioural results of Experiment 2. Panel A shows the angular error for the different combinations of the feature cue (selective vs. neutral) and task cue (selective vs. neutral). The orange-coloured dots depict the mean angular error of each participant for selective task cues and the green coloured dots for the neutral task cue condition. The vertical line indicates the condition average. Panel B represents the correct response rate in the recognition task. Panel C shows the time required to initiate the mouse movement and Panel D the response times in the recognition task.

For the recognition task, the percentage of correct responses (see figure 3B) was used to assess working memory accuracy. Here, selective feature cues increased performance, *F*(1,21) = 27.46, *p* < .001, *p_crit_* = .007, *η^2^_p_* = 0.57. As in the continuous task, the task cues had no effect on accuracy, *F*(1,21) = 0.36, *p* = .554, *p_crit_* = .043, *η^2^_p_* = 0.02. In general, performance was better for location than for orientation recognition, *F*(1,21) = 24.97, *p* < .001, *p_crit_* = .014, *η^2^_p_* = 0.54.

As in Experiment 1, time to mouse movement onset (see figure 3C) was utilized as a measure for speed of response initiation in the continuous report task. Response initiation did not differ between location and orientation adjustment trials, *F*(1,21) = 1.99, *p* = .173, *p_crit_* = .027, *η^2^_p_* = 0.09. Responses were speeded by a selective feature cue, *F*(1,21) = 48.77, *p* < .001, *p_crit_* = .007, *η^2^_p_* = 0.70, as well as by a selective task cue, *F*(1,21) = 47.66, *p* < .001, *p_crit_* = .014, *η^2^_p_* = 0.69. After correcting for hidden multiplicity within the ANOVA none of the possible interactions remained significant (all p-values > .027, corresponding critical p-values = .014).

Responses to the recognition task were faster after selectively cueing either of the features, *F*(1,21) = 36.69, *p* < .001, *p_crit_* = .014, *η^2^_p_* = 0.64. The selective task cue also decreased response times, *F*(1,21) = 27.79, *p* < .001, *p_crit_* = .021, *η^2^_p_* = 0.57. Response times in general were faster for location than for orientation recognition, *F*(1,21) = 55.17, *p* < .001, *p_crit_* = .007, *η^2^_p_* = 0.72. Finally, the effect of the feature cue differed between the cued features, *F*(1,21) = 7.93, *p* = .010, *p_crit_* = .029, *η^2^_p_* = 0.27, in the way that selective feature cues decreased response times more strongly in location, *t*(21) = -5.44, *p_adj_* < .001, *d_z_* = -1.16, CI 95% [-205.39 -91.84], than orientation recognition trials, *t*(21) = -3.77, *p_adj_* = .001, *d_z_* = -0.80, CI 95% [-103.04 -29.73].

Summarized, also Experiment 2 indicated an overall performance benefit of selective feature cues. While the task cues did not consistently affect performance, a benefit was shown regarding the time required for response initiation in both the continuous report task and the recognition task. This is especially true for a continuous report of orientation, since here a selective feature cue further amplified the acceleration of response initiation following a selective task cue.

### 3.2. EEG results

#### 3.2.1. Experiment 1

##### 3.2.1.1. Channel-based analysis

Analyses on EEG level were focused on the left (PO3, PO7, PPO5h, P7, P5) and right (PO4, PO8, PPO6h, P8, P6) posterior electrodes as well as on the left (CP3, CCP5h, CCP3h, C3) and right (CP4, CCP6h, CCP4h, C4) electrodes over sensorimotor sites. In a first step, time-frequency data were collapsed over all electrode clusters and selective and neutral trials were contrasted by a cluster-based permutation procedure, which revealed a broad significant difference in mu and beta frequency ranges following the feature cue (see figure 4A). Based on this time-frequency cluster, a rm-ANOVA was run with the factors *condition* (location vs. orientation vs. neutral feature cue) x *hemisphere* (left vs. right) x *caudality* (central vs. posterior). Importantly, this ANOVA revealed a 3-way interaction, *F*(2,46) = 3.45, *p* = .040, *η^2^_p_* = 0.13 (see figure 4 C-F).

**Figure 4.**
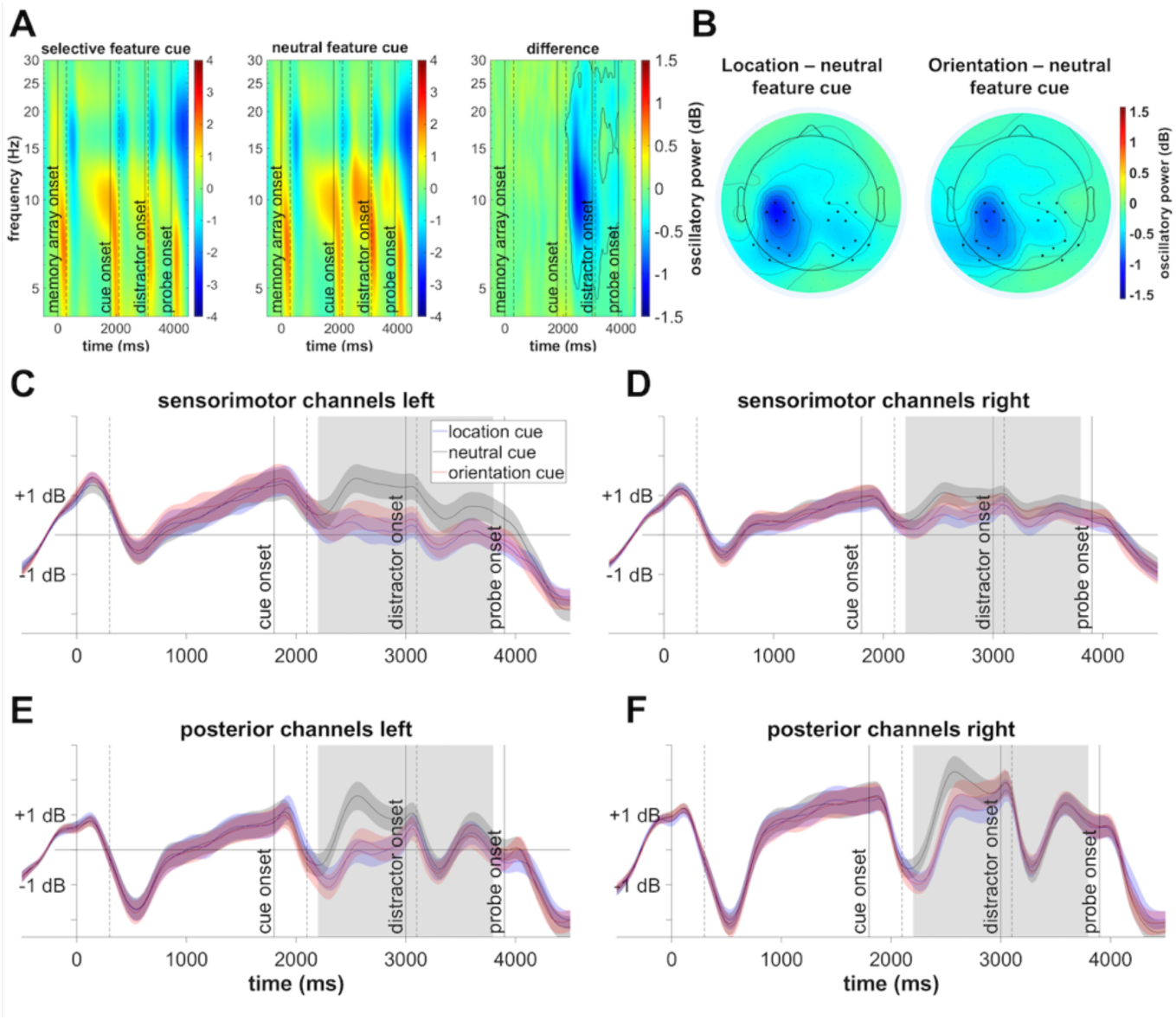
Results of the channel-based analyses. Panel A shows the time-frequency distribution of the activity averaged across channels and subjects for the selective cue, the neutral cue condition, and the difference between selective and neutral activity. The vertical solid lines indicate relevant events throughout the trial (0 ms - memory array onset, 1800 ms - selective cue onset, 3000 ms - visual distractor onset, 3900 - ms memory probe onset). Stimulus offset times are indicated by dashed vertical lines. Panel B illustrates the topographical distribution of the location-minus-neutral condition activity (left) and the orientation-minus-neutral condition activity (right) averaged across the significant cluster (see solid line in panel A). Panel C-F highlights the time course of oscillatory power in the significant frequency range (8-15 Hz) for the four electrode clusters of interest. The standard error of the mean is indicated by shaded areas surrounding the condition average. The grey areas depict significant time windows obtained by conducting the cluster-based permutation analysis (2200 - 3300 ms; see panel A).

**Figure 5.**
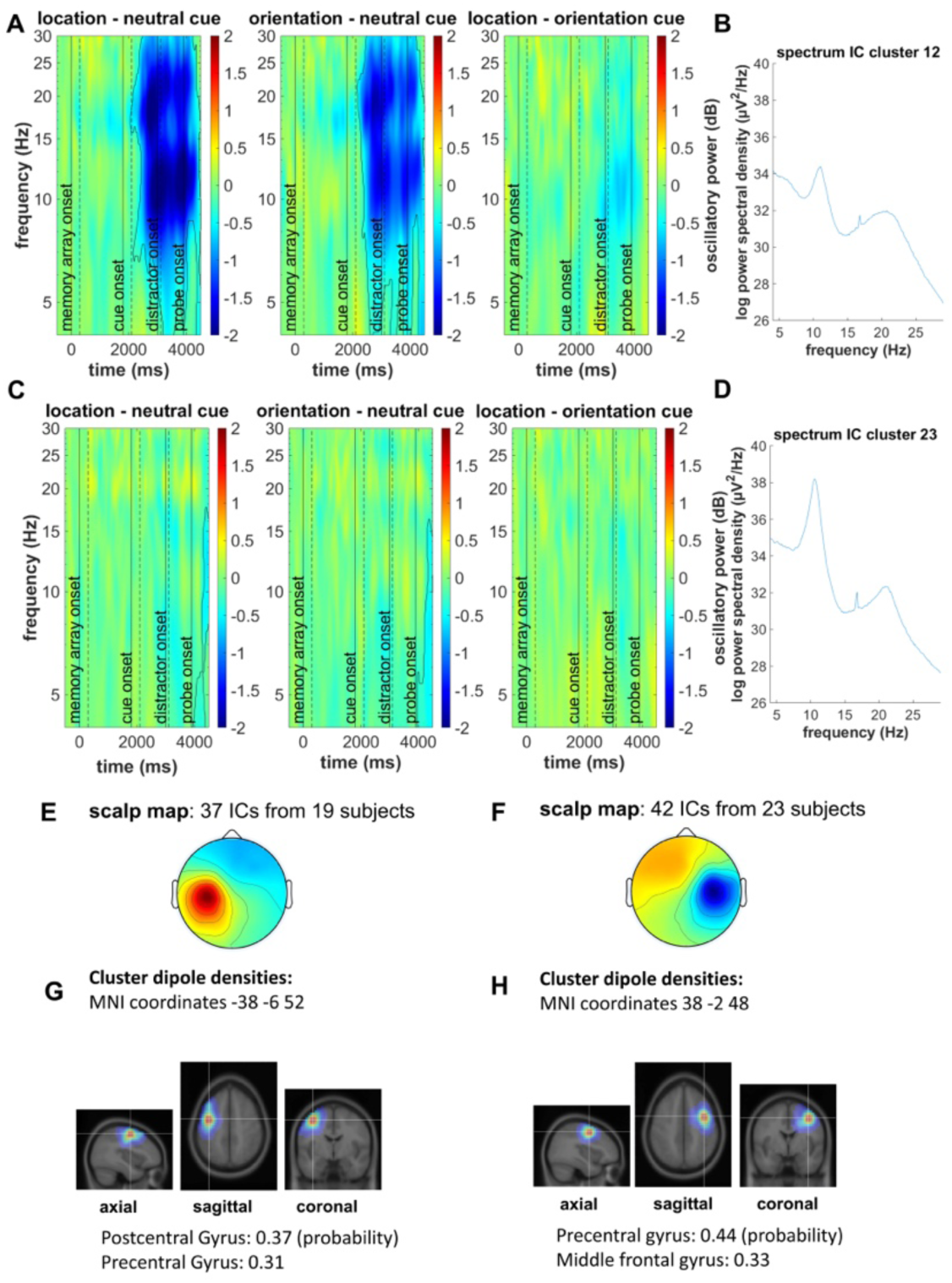
Feature cue effects for contralateral vs. ipsilateral mu/beta IC clusters (Experiment 1). Panel A depicts a time-frequency plot showing the condition differences contralateral to the responding hand. Time-frequency clusters differing between the conditions are indicated by a solid (-) line. Stimulus offset times are indicated by dashed vertical lines. Panel B shows the spectral power of the left hemispheric IC cluster. Panel C and D depict the respective results for the right hemispheric IC cluster. Panel E and G show the scalp map and the respective dipole density distribution of the left hemispheric IC cluster. Panel F and H depict the scalp map and dipole density distribution for the right hemispheric IC cluster.

When only considering the posterior recording sites, there was a *condition x hemisphere* interaction, *F*(2,46) = 9.37, *p* = .001, *p_crit_* = .042, *η^2^_p_* = 0.28, ε = .80. The condition effect was stronger over left posterior sites, *F*(2,46) = 21.67, *p* = .001, *p_crit_* = .017, *η^2^_p_* = 0.29, ε = .69, than over right posterior sites, *F*(2,46) = 8.53 *p* = .003, *p_crit_* = .05, *η^2^_p_* = 0.27, ε = 0.73. The 3-way interaction can be explained by the fact that for the central recording sites, there was an even stronger *condition x hemisphere* interaction, *F*(2,46) = 11.09, *p* < .001, *p_crit_* = .042, *η^2^_p_* = 0.33, χ = 0.79. Again, the feature cue effect was stronger over the left hemisphere, *F*(2,46) = 31.74, *p* < .001, *p_crit_* = .013, *η^2^_p_* = 0.58, χ = 0.69, than over the right hemisphere, *F*(2,46) = 10.61, *p* < .001, *p_crit_* = .038, *η^2^_p_* = 0.32, χ = 0.87.

Summarized, there was a stronger hemispheric difference regarding the feature cue effect (selective vs. neutral) over the sensorimotor cortex than over posterior visual areas. This might indicate that the lateralized mu/beta effect was associated with the planning of the right-handed motor response following selective feature cues. To further support this assumption, we looked specifically at oscillatory power from estimated neural sources in left vs. right sensorimotor cortex based on an IC-clustering approach.

##### 3.2.1.2. IC-based analysis

We observed two IC clusters that featured the typical characteristics of the mu and beta oscillatory response in preparation for responses and during their execution (see Jenson et al., 2019; Jenson & Saltuklaroglu, 2021; Schneider et al., 2017). These characteristics include a spectral response (FFT) with a peak in both the alpha (8-14 Hz) and beta (15-30 Hz) frequency range (see figures 5 B & D, 8 B & D), an average dipole location within left- or right-hemispheric pre-motor or motor cortex, and a scalp distribution of the IC cluster with strongest activity over left- or right motor areas (see figures 5 E-H, 7 B, C, E & F).

Dipole density analysis indicated estimated neural sources in the left sensorimotor cortex for IC cluster 12 (located with 37% probability in the precentral gyrus and 31% probability in the post central gyrus; see figure 5C) and right sensorimotor cortex for IC cluster 23 (located with 44% probability in the precentral gyrus and 33% probability in the middle frontal gyrus; see figure 5F).

All conditions were contrasted by separate cluster-based permutation procedures. Both the location and the orientation feature cues led to differences in oscillatory power in mu and beta frequency range relative to the neutral cue condition (see figure 5 A). Importantly, only for the left (contralateral) IC cluster, this effect appeared clearly prior to the onset of the memory probes demanding the actual motor response. For the right (ipsilateral) IC cluster, the minor differences between the selective and neutral cue conditions were only evident after memory probe presentation (see figure 5 C).

#### 3.2.1. Experiment 2

##### 3.2.1.1. Channel-based analysis

Oscillatory power averaged over the four electrode clusters was compared between the selective and neutral feature cue conditions and revealed a time-frequency cluster with a significant difference after the feature cue presentation (see figure 6). Like for Experiment 1, the subsequent rm-ANOVA revealed a significant *condition x hemisphere x caudality* interaction, *F*(2,44) = 4.47, *p* = .017, *η^2^_p_* = 0.17. While a subsequent ANOVA on posterior channels revealed no hemispheric difference regarding the condition effect, *F*(2,44) = 0.02, *p*= .982, *p_crit_* = .05, *η^2^_p_* < 0.01, the same analysis focusing on the two central electrode clusters found a significant *condition x hemisphere* interaction, *F*(2,44) = 4.47, *p < .*001*, p_crit_ = .*025*, η^2^_p_* = 0.27. This again suggested that the stronger suppression of mu and beta power following selective feature cues was related to motor preparation processes.

**Figure 6.**
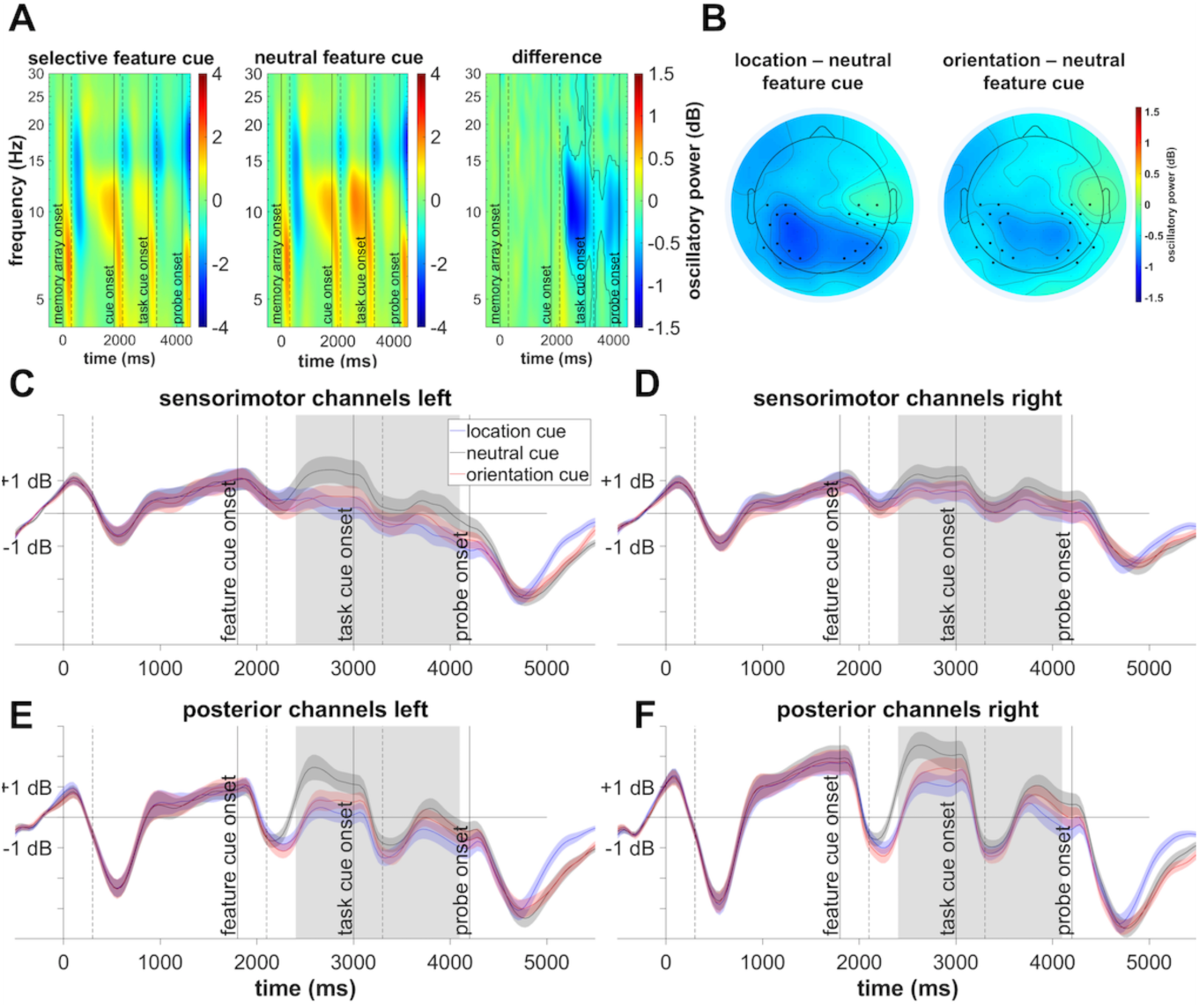
Results of the channel-based analyses (Experiment 2). Panel A shows the time-frequency distribution of the activity averaged across channels and subjects for the selective cue, neutral cue condition, and the difference between the two conditions. The vertical solid lines indicate relevant events throughout the trial (0 ms –memory array onset, 1800 ms – selective cue onset, 3000 ms – task cue onset, 4200 memory probe onset) and the solid line (-) marks the cluster in which the conditions differ significantly. Stimulus offset times are indicated by dashed vertical lines. Panel B illustrates the topographical distribution of the location-minus-neutral condition activity (left) and the orientation-minus-neutral condition activity (right averaged across the significant cluster (see panel A)). Panel C-F highlights the time course of the significant frequency range for the four electrode clusters of interest. The standard error of the mean is indicated by the shaded area surrounding each condition average. The grey areas depict significant time windows (2200 ms - 4100 ms) obtained from the cluster-based permutation analysis (see panel A).

**Figure 7.**
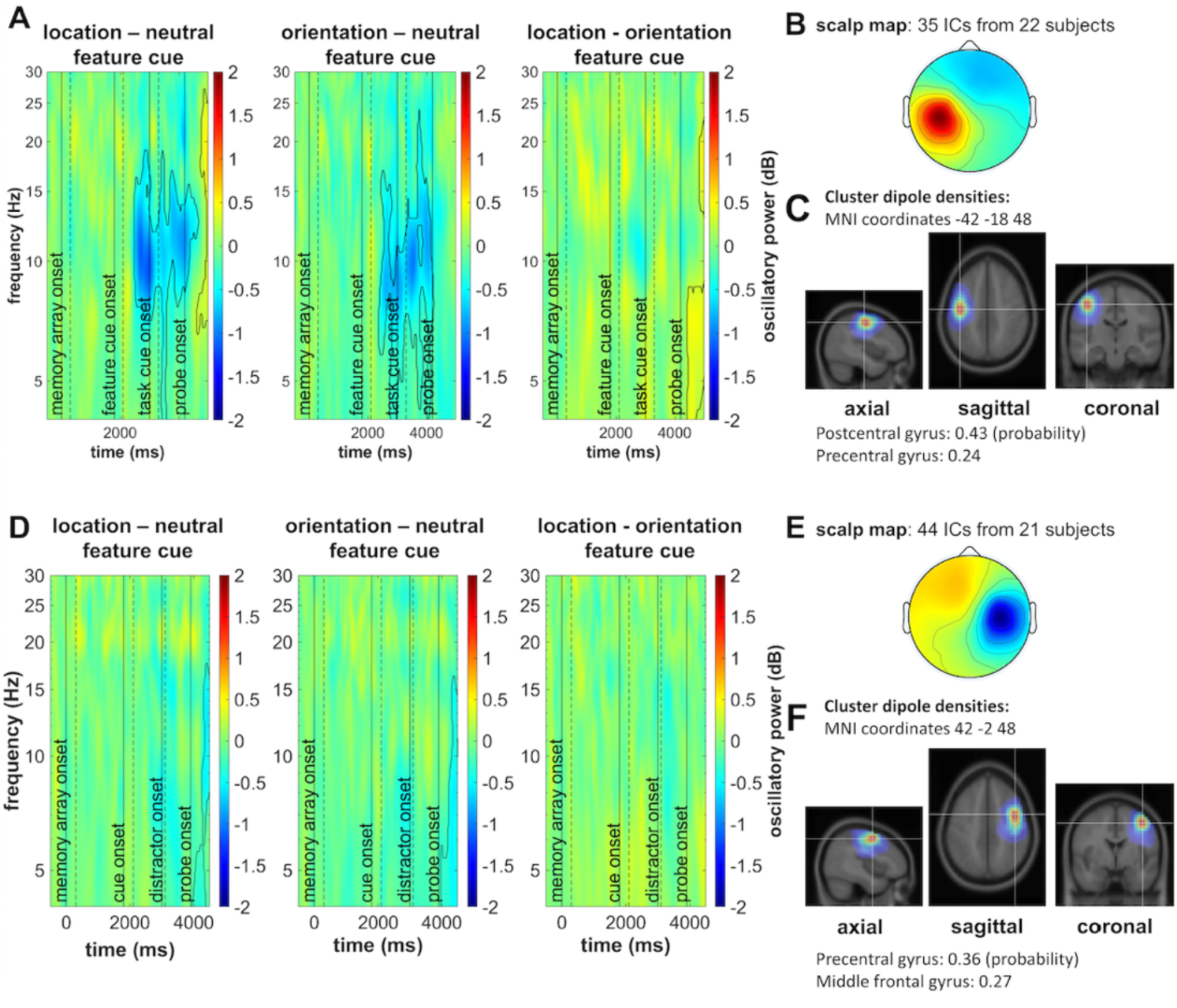
Feature cue effects for contralateral vs. ipsilateral mu/beta IC clusters (Experiment 2). Panel A depicts the differences in oscillatory power between conditions for all pair-wise contrasts between the feature cues at the contralateral IC cluster. Time-frequency clusters differing between the conditions are indicated by a solid (-) line. Panel B shows the scalp map of the left-hemispheric IC cluster and Panel C the respective dipole densities. Panel D-F depict the same results for the right sensorimotor cortex.

**Figure 8.**
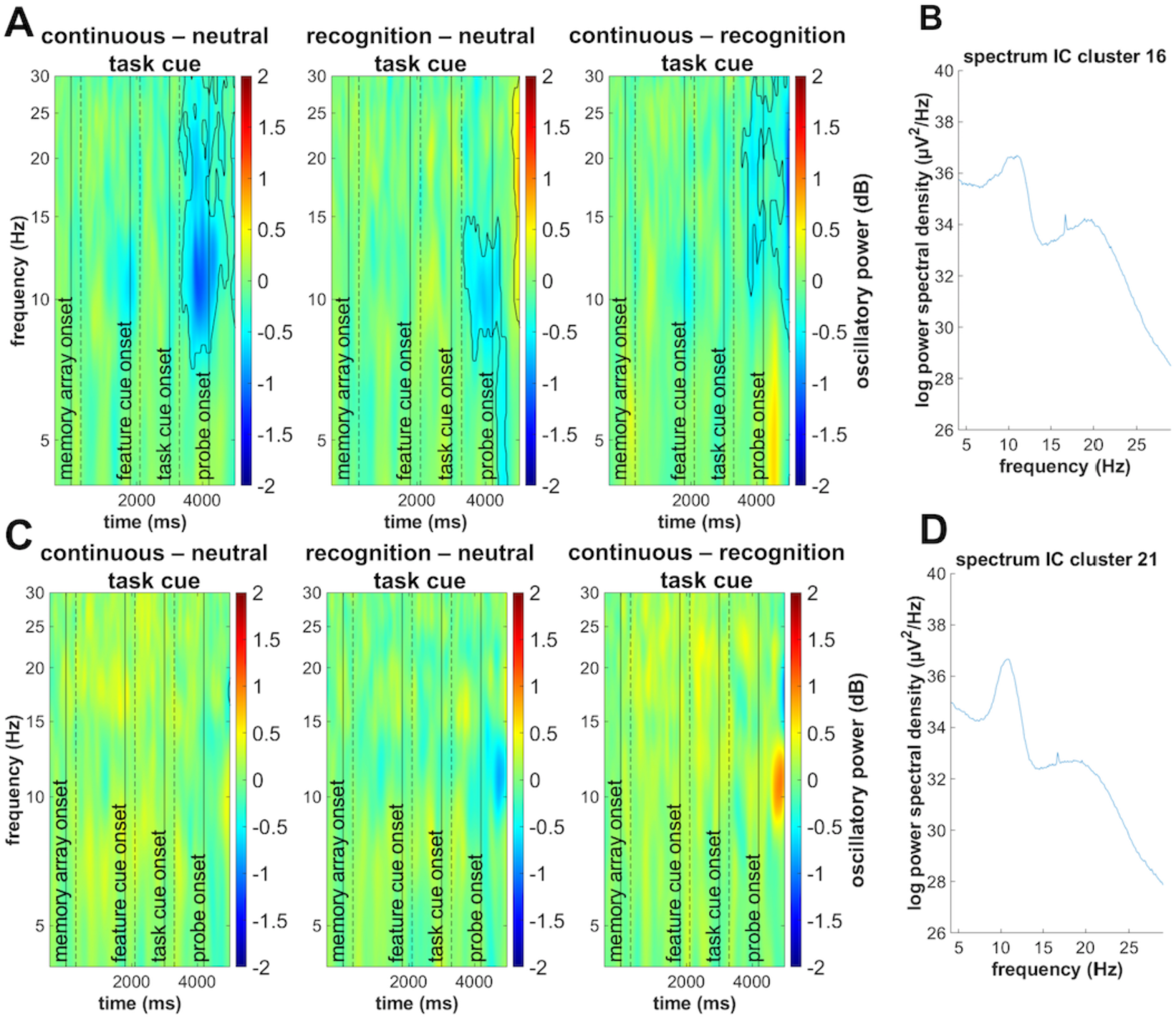
Task cue effects for contralateral vs. ipsilateral mu/beta IC clusters (Experiment 2). Panel A depicts the difference in oscillatory power for all pairwise contrasts between the three task cues for the left-hemispheric IC cluster. Time-frequency clusters differing between the conditions are indicated by a solid (-) line. Panel B shows the spectral power for the left-hemispheric IC cluster. Panel C and D depict the same results for the right-hemispheric IC cluster.

##### 3.2.1.2. IC-based analysis

As revealed by the dipole density analysis, the left-hemispheric mu/beta IC cluster 16 (precentral gyrus: 24%; postcentral gyrus: 43%) and the right-hemispheric mu/beta IC cluster 21 (precentral gyrus: 36%; middle frontal gyrus: 27%) were estimated in sensorimotor and premotor cortex with a high probability. Only for the left-hemispheric mu/beta IC cluster, the cluster-based permutation procedure revealed a stronger suppression of oscillatory power following the location cue than following the neutral cue. Similarly, there was a stronger suppression of oscillatory power after the orientation cue than after the neutral cue. The two selective feature cue conditions (location vs. orientation) revealed no difference in this regard (see figure 7A). This highlights that, even when the to-be-executed task was not yet fully specified, cuing of the target feature resulted in a stronger suppression of mu and beta oscillatory power, but only in the sensorimotor cortex contralateral to the future response.

Comparing the data based on the second task cue (see figure 8) revealed a significant cluster of stronger mu and beta suppression following a continuous report task cue than a neutral task cue. Cuing a recognition task resulted in a stronger suppression of mu oscillatory power compared to a neutral task cue. There was also a difference between the selective task cues. The continuous report task cue resulted in a stronger mu and beta suppression starting at around 748 ms after cue onset. We further analysed whether the difference in oscillatory power between selective (continuous report or recognition) and neutral task cues differed as a function of feature cue type. The time-frequency window for this analysis resulted from the comparison of the combined selective task cues and the neutral task cue condition with the cluster-based permutation procedure described above. This was done to test whether the differences in the oscillatory response between the task cue conditions were actually based on the process of selecting a specific feature that was only postponed to a time when the actual task had been specified. These post-hoc analyses could not reveal a difference in the task cue effect depending on the prior feature cues, *F*(4,84) = 1.50, *p* = .210, *p_crit_* = .05, *η^2^_p_* = 0.07, making it unlikely that delayed feature selection can fully explain the effects of the task cue. However, we cannot completely rule out this alternative explanation, since there was no longer a reliable effect between selective and neutral task cues when only considering trials with neutral feature cues (continuous report: *t*(21) = -1.98, *p_adj_* = .122, *d_z_* = -0.42, CI [-0-78 0.02]; recognition task: *t*(21) = -1.26, *p_adj_* = .221, *d_z_* = -0.23, CI [-0.53 0.13]).

When comparing the oscillatory response of the contralateral mu/beta cluster between trials with fast vs. slow responses based on a median split within each experimental condition, we found a stronger suppression of oscillatory power in the mu frequency range to be linked to fast responses. A post-hoc ANOVA based on the significant time-frequency cluster revealed that this effect appeared independent of experimental conditions. There was no interaction of the RT effect with the type of feature cue, *F*(2,42) = 0.74, *p* = .483, *p_crit_* = .043, *η^2^_p_* = 0.03, and it did not differ between the continuous report and the recognition task, *F*(2,42) = 0.69, *p* = .507, *p_crit_* = .05, *η^2^_p_* = 0.03. Also, the three-way interaction was non-significant, *F*(4,84) = 2.247, *p* = .071, *p_crit_* = .029, *η^2^_p_* = 0.10.

## 4. Discussion

By means of oscillatory parameters of the EEG, the present study demonstrates that the selection of visual features stored in working memory results in the concurrent selection of preparatory motor codes. This is true even when an individual feature of a single item stored in working memory is selected and when the exact, to-be-executed task is still unknown.

In line with earlier research on feature selection in working memory (Hajonides et al., 2020; Niklaus et al., 2017; Sasin & Fougnie, 2020; Ye et al., 2016), we could show in Experiment 1 that a retrospective cue towards a single feature of a visual object caused a performance benefit relative to a neutral cue condition, both in terms of working memory precision and the speed of response initiation (see figure 2). On the electrophysiological level, we observed a suppression of alpha (or mu) and beta power (∼8 – 30 Hz) at posterior and centro-parietal recording sites that was stronger following selective feature cues (see figure 4). Importantly, and in line with a prior investigation from our lab (see Schneider et al., 2017), the centro-parietal electrode clusters featured an increased hemispheric difference, with a stronger suppression of oscillatory power over the left-hemispheric electrodes. In addition to the suppression of posterior alpha power already shown in the context of feature selection in working memory (Hajonides et al., 2020), this indicates an oscillatory effect prior to memory probe presentation that appeared specifically over the motor cortex contralateral to the responding hand. Earlier research has shown that this contralateral suppression of oscillatory power appeared prior to both left-sided and right-sided motor responses (Schneider et al., 2020; van Ede et al., 2019; Zickerick et al., 2021).

The IC-clustering approach supported the contralateral mu and beta suppression effect (see figures 5B). We obtained both a left-hemispheric and right-hemispheric IC cluster with the typical spectral peaks in mu and beta frequencies and estimated neural sources in sensorimotor and pre-motor cortex. Importantly, only the left-hemispheric cluster (i.e., the cluster contralateral to the executed motor response) featured a stronger suppression of mu and beta power following the selective feature cues. The ipsilateral mu/beta IC cluster showed differences in oscillatory power only after memory probe presentation (see figure 5E). This oscillatory pattern has typically been linked to the planning of a response prior to its actual execution, both when selecting information relevant for later action from working memory (e.g. Boettcher et al., 2021, S. 2; Schneider et al., 2017, 2020; van Ede, 2018) and prior to self-paced movements (Leocani et al., 1997; Pfurtscheller et al., 2000; Zhuang et al., 1997). Thus, whereas the posterior suppression of alpha power can be associated with the selection of a visuospatial representation stored in working memory, the suppression in mu and beta frequencies over contralateral centro-parietal areas can be linked to the concurrent selection of a motor code associated with the cued visual feature. The stronger suppression of mu and beta power following the selective retro-cues appeared both when location and when orientation was the relevant feature. This clearly shows that such motor-related processing is not specific to the selection of individual items from working memory (Boettcher et al., 2021; Schneider et al., 2017; van Ede et al., 2019).

The second experiment focused on the question whether the precise definition of the upcoming task is a prerequisite for the development of a motor representation in working memory. Olivers and Roelfsema (2020) proposed that the difference between attended and non-attended information stored in working memory might be the coupling of a sensory representation with an action plan. However, it remains unclear whether this coupling requires the precise knowledge of the action to be executed next or whether it also happens when the to-be-executed action is still uncertain. In the latter case, a prospective motor code would be created for dealing with the continuous report task (see Experiment 1), even though it would not be necessary for the visual comparison of the memory probe with the stored information (recognition task).

The current study strongly speaks in favour of this latter assumption. Analogous to Experiment 1, the channel-based analysis revealed a stronger suppression of mu and beta power at posterior and centro-parietal sensors following the selective cuing of the location or orientation of the stored visual object (i.e., following the first retro-cue). This effect was stronger at electrodes contralateral to the side of the to-be-executed actions, but only for the centro-parietal electrode clusters over left and right sensorimotor cortex (see figure 6).

Moreover, the IC clustering approach strengthened our assumption that this oscillatory effect was linked to motor-related processes and not simply related to feature selection effects reflected in oscillatory patterns in more posterior brain regions. Again, the left-hemispheric (contralateral) and right-hemispheric (ipsilateral) IC cluster featured the typical double spectral peaks in mu and beta frequency ranges and showed the highest estimated dipole density in sensorimotor and pre-motor cortex. Comparable to Experiment 1, only the contralateral cluster revealed a stronger suppression of mu and beta power following the selective feature cues. Thus, Experiment 2 shows that a motor-related code was selected at a time when it was not yet clear whether it would be required for the next action to be performed. The relevance of this oscillatory effect for behavioural performance was indicated by a comparison of fast vs. slow response trials within the experimental conditions. Greater suppression of mu oscillatory power prior to the presentation of the memory probe display was associated with fast responses (see figure 9) and this effect only occurred for the contralateral mu/beta IC cluster.

**Figure 9.**
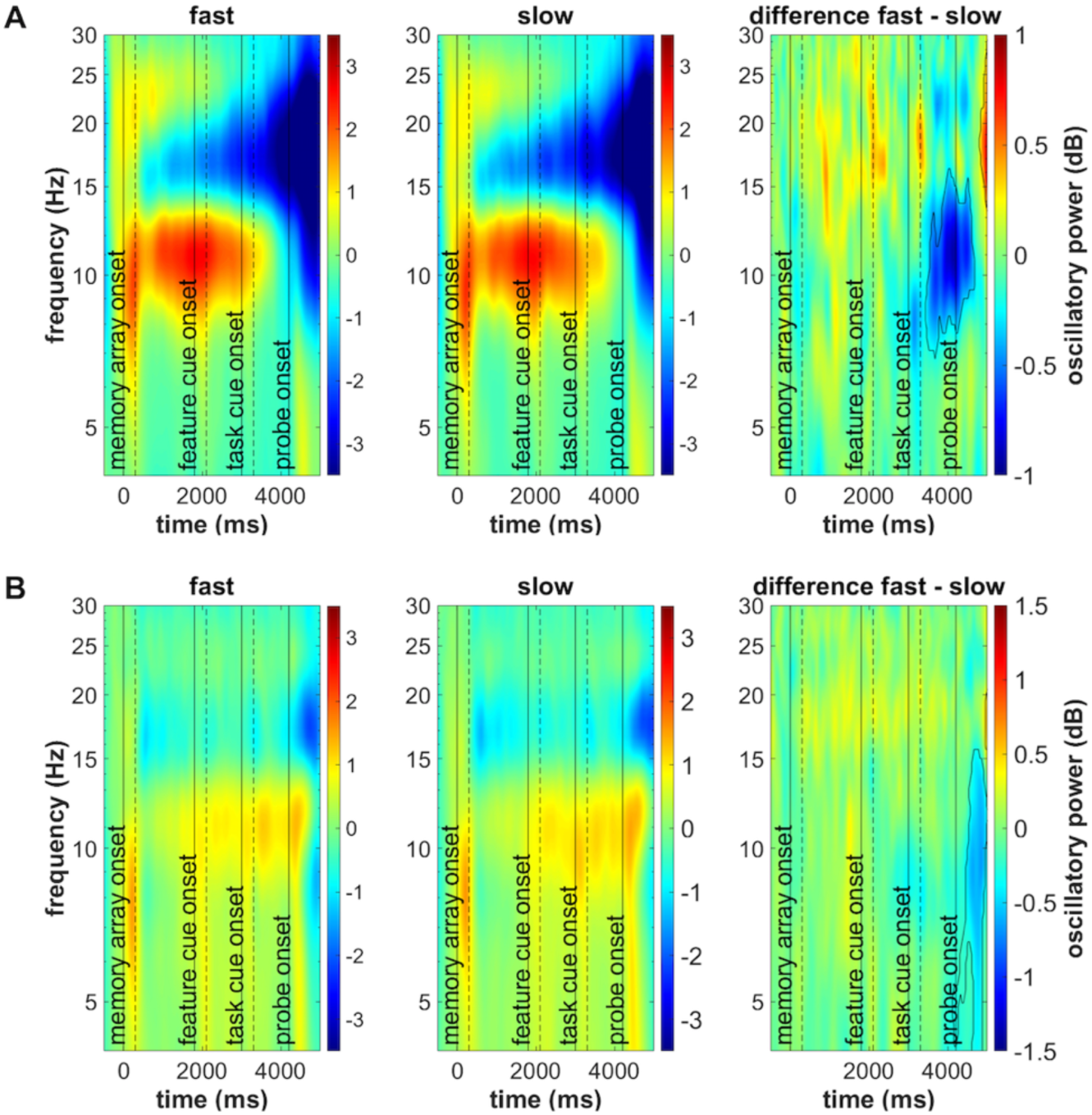
Response time effects for contralateral vs. ipsilateral mu/beta IC clusters (Experiment 2). Panel A depicts the oscillatory power of fast and slow trials and their difference for the left-hemispheric IC cluster. The time-frequency clusters differing between the conditions are indicated by a solid (-) line. Panel B shows the same for the right-hemispheric IC cluster.

In line with these findings, Henderson, Rademaker and Serences (2021) proposed that information stored in working memory can be represented in a rather flexible format, based on the requirements of the task at hand. For example, when it is required to compare the representation of a visual stimulus to a memory probe, the stored mental representation might be based on a “retrospective” visual code. On the other hand, when the information stored in working memory is used to precisely manipulate an object by means of a goal-directed movement, a prospective motor code might additionally be required. This is in line with the pattern of oscillatory activity following the task cues. The contralateral mu/beta IC cluster featured a stronger suppression of oscillatory power following a selective task cue than following a neutral cue (i.e., no task definition until memory probe presentation). This pattern, however, differed between the different task cues. Following the continuous report task cue, we observed a stronger suppression of both mu and beta frequencies (see figure 8A), whereas this modulation of oscillatory power was limited to the mu frequency range following the recognition task cue. In contrast to the continuous report task where the participant has all the necessary information in working memory to respond as soon as the memory probe appears, one must consider new information (the probe stimulus) and compare it to the contents of working memory in the recognition task. Therefore, it is not possible to respond immediately, but only after the memory probe has been compared to the relevant information stored in working memory. It is interesting that there is still some motor preparatory activity (see figure 8A; recognition vs. neutral task cue) in this case, which further strengthens the notion of a motor planning process even when the actual response to be executed is still ambiguous (see also: Nasrawi & van Ede, 2021).

Further, when comparing the two selective task cue conditions directly, the contralateral IC cluster revealed a stronger suppression of mu and beta oscillatory power prior to a cued continuous report task. This suggests that subjects potentially relied more strongly on a prospective motor code when preparing for the continuous report task. Alternatively, subjects might have been preparing for two different tasks in terms of the mere motor requirements.

While the continuous report task required an arm movement (in order to move the computer mouse), the recognition task required simply a key press (i.e., a finger movement).

Furthermore, it has to be considered that the comparison of the two selective task cue conditions might be confounded by task difficulty. In this sense, different oscillatory patterns of the contralateral mu/beta IC cluster between the continuous report and recognition task would not be directly based on differences in the type of mental representation required (prospective vs. retrospective mental representations). Rather, the preparation for a generally more difficult task could be associated with the fact that motor planning processes become necessary to a greater extent even before memory probe presentation. Based on the current experiment, we cannot distinguish between these alternative explanations. To allow for a more specific conclusion in this regard, the two tasks need to be linked to identical motor responses and matched in terms of difficulty. Nonetheless, the present data clearly show that compared to the neutral condition, both selective cues brought about a further specification of the mental representation of the to-be-executed task.

### 4.1. Conclusion

The current study sheds light on the role of motor representations for the goal-oriented processing of information in working memory. The first experiment showed that the retrospective selection of object features can entail the selection of corresponding motor representations. This shows that individual features of a visual object can also be stored and selected as a visual (input) and motor (output) code. The second experiment further highlighted that such motor codes are prospectively generated even when the exact to-be-performed task is not fully specified. Thus, it can be assumed that the observed effects in the mu and beta frequency range reflect a higher-level representational state of the stored working memory content that serves as a basis for later response planning processes once the to-be-executed task was cued. This shows that working memory flexibly stores different kinds of memory representations in such a way that the best possible conditions for the execution of a given task are provided.

## 5. Acknowledgements

We would like to thank Tobias Blanke for programming the experiment.

## 6. Data and code availability statement

All relevant data (including the raw EEG data and the scripts used for the analysis) will be made publicly available at the OSF after acceptance for publication.

## 9. Supplementary

**Supplementary A.**
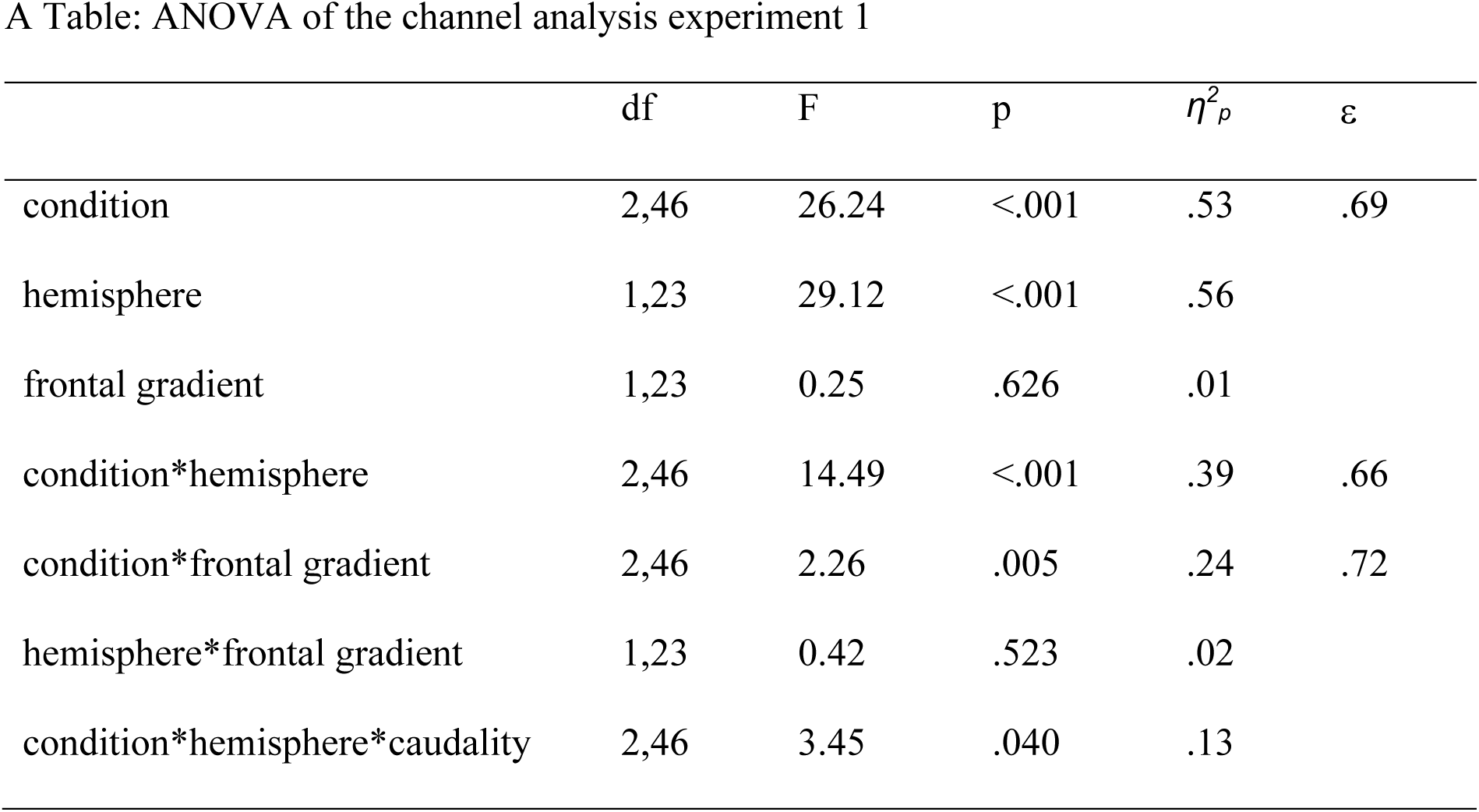
3-factor ANOVA for experiment 1: condition (location vs. orientation vs. neutral feature cue) x hemisphere (left vs. right hemisphere) x caudality (central vs. posterior channels).

**Supplementary B.**
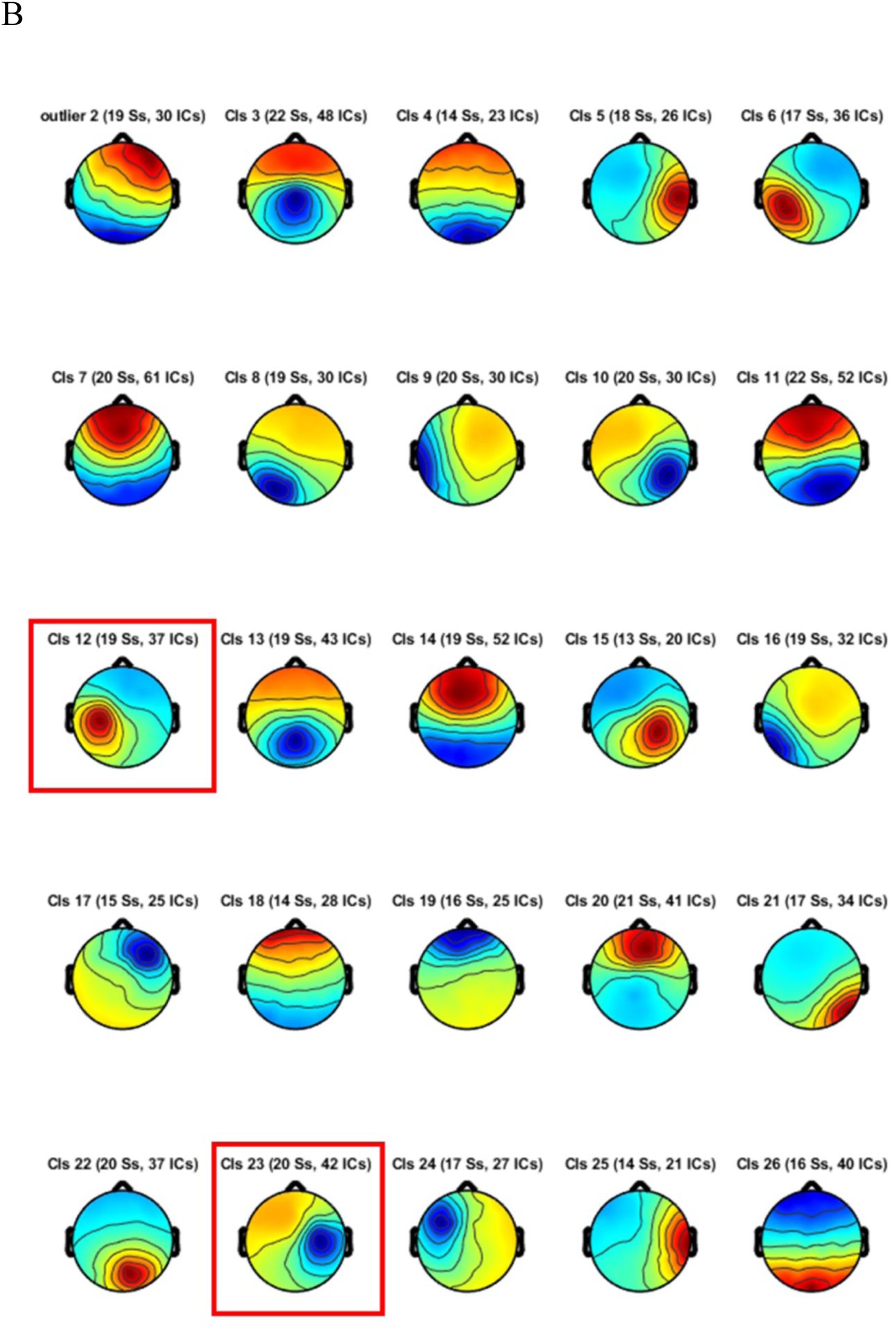
Scalp topographies of independent component clusters in Experiment. Mean topographies of 24 IC clusters. The clusters representing left hemispheric and right-hemispheric mu/beta oscillatory activity are marked in red. The number of subjects (Ss) and ICs contributing to each cluster are given above each topography. In addition to the 24 clusters, the mean topography for the outlier cluster is provided in the top-left corner.

**Supplementary C.**
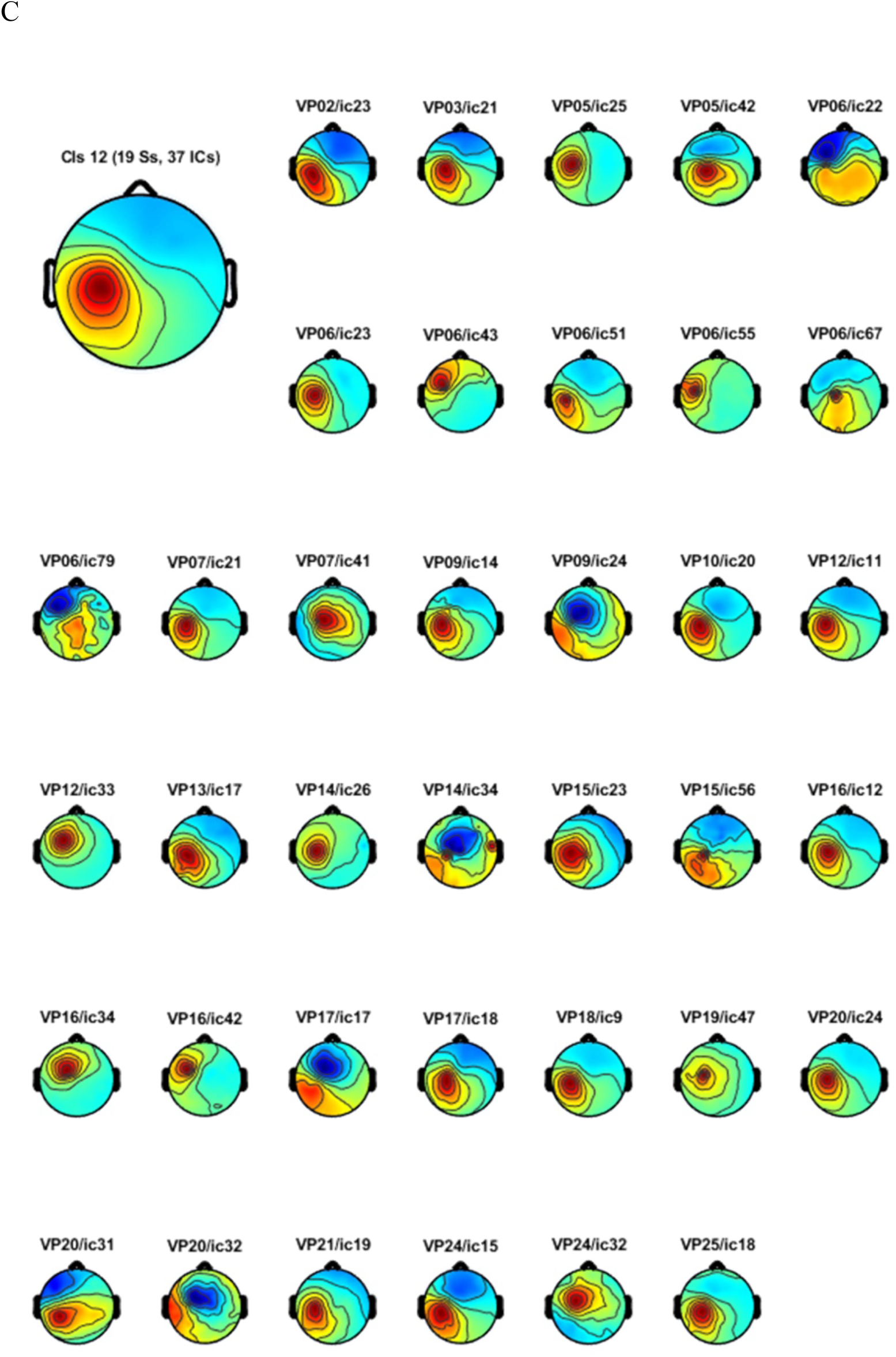
Individual scalp topographies contributing to the contralateral IC cluster in experiment 1.

**Supplementary D.**
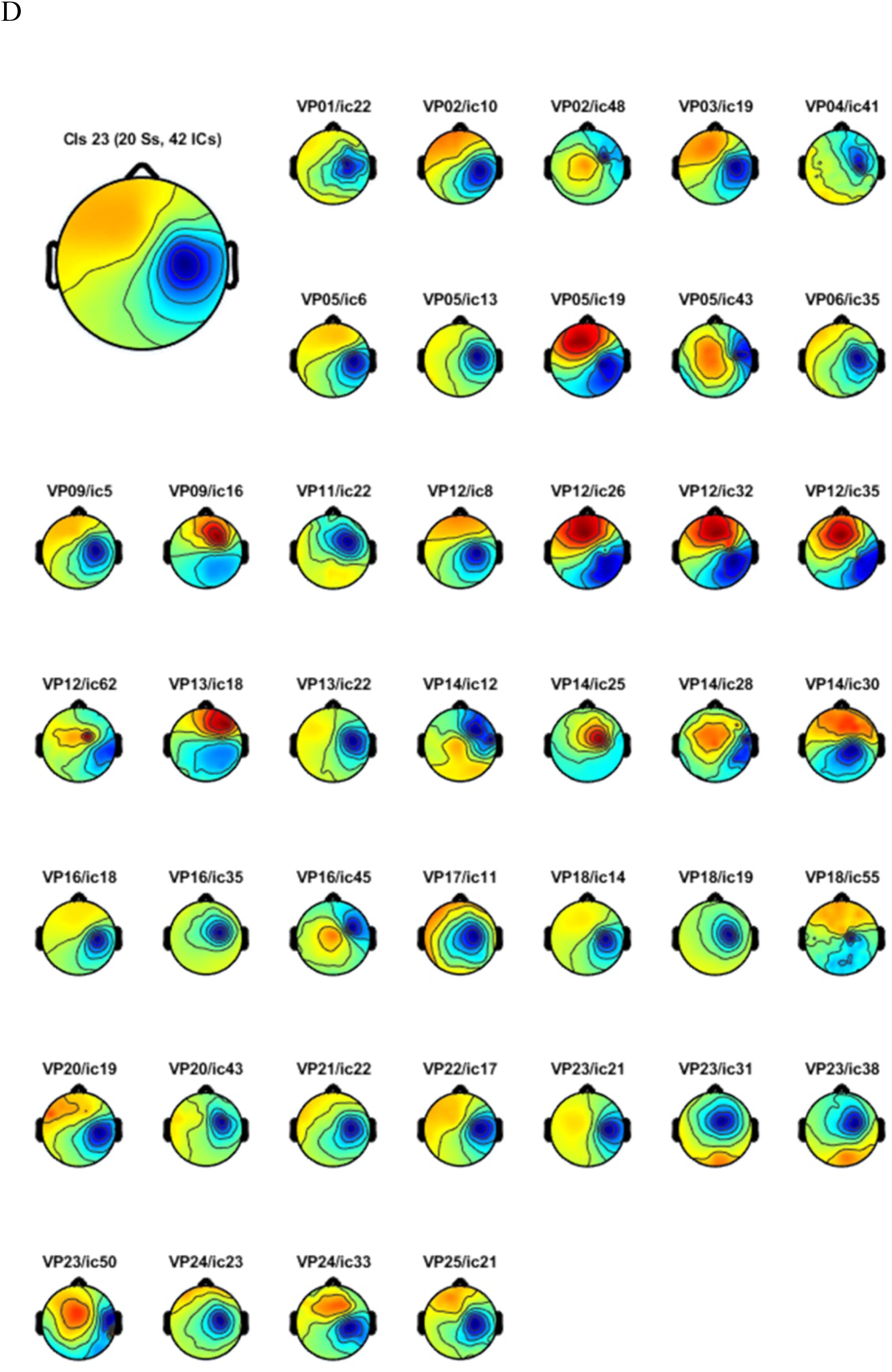
Individual scalp topographies contributing to the ipsilateral IC cluster in experiment 1.

**Supplementary E.** 3-factor ANOVA for experiment 2: condition (location vs. orientation vs. neutral feature cue) x hemisphere (left vs. right hemisphere) x caudality (central vs. posterior channels).

| | df | F | p | $\eta^2_p$ |
| --- | --- | --- | --- | --- |
| condition | 2,44 | 13.55 | <.001 | .38 |
| hemisphere | 1,22 | 17.27 | <.001 | .44 |
| frontal gradient | 1,22 | 0.68 | .419 | .03 |
| condition*hemisphere | 2,44 | 3.63 | .035 | .14 |
| condition*frontal gradient | 2,44 | 4.34 | .019 | .17 |
| hemisphere*frontal gradient | 1,22 | 0.31 | .581 | .01 |
| condition*hemisphere*caudality | 2,44 | 4.47 | .017 | .17 |

**Supplementary F.**
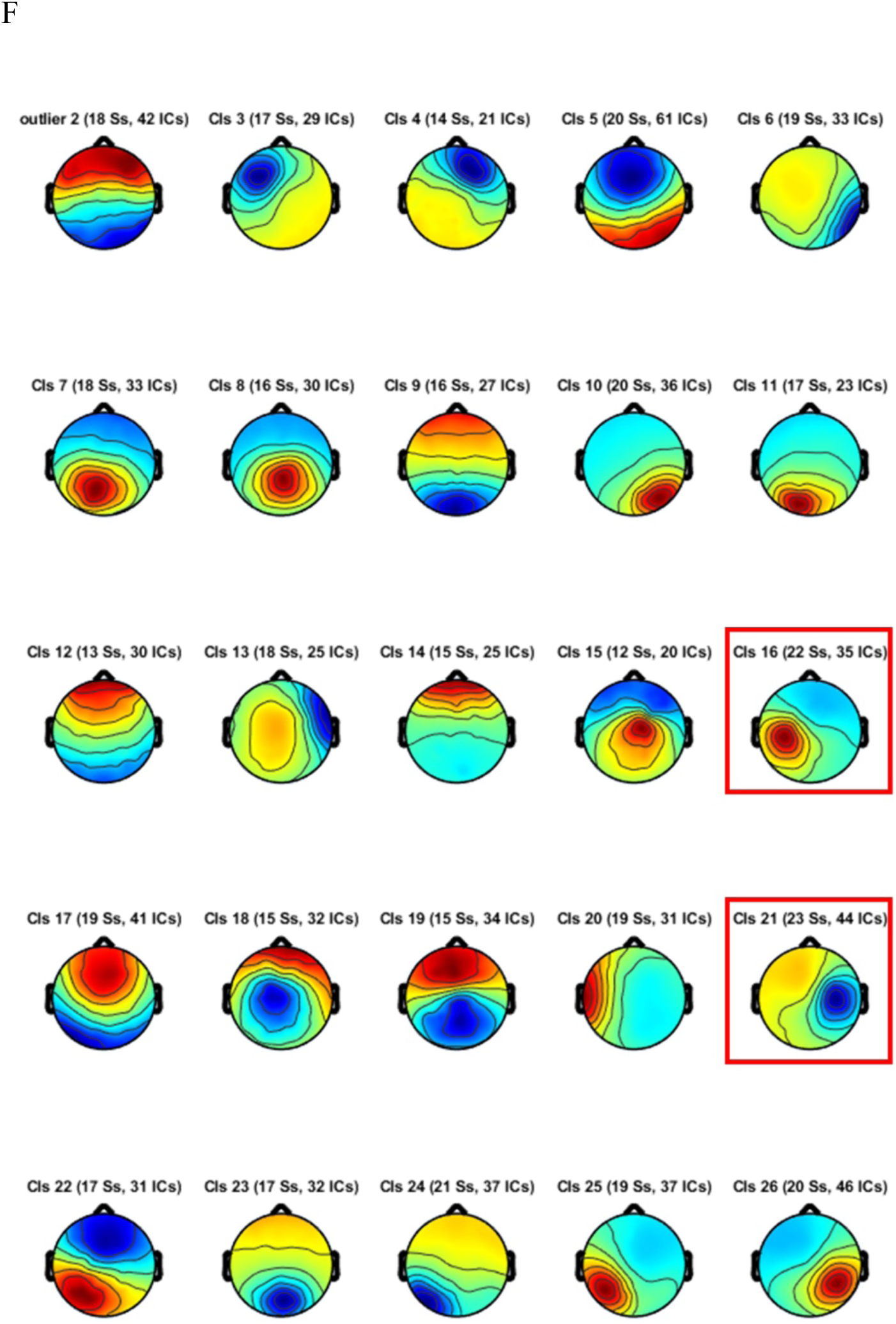
Scalp topographies of independent component clusters in Experiment. Mean topographies of 24 IC clusters. The clusters representing left hemispheric and right-hemispheric mu/beta oscillatory activity are marked in red. The number of subjects (Ss) and ICs contributing to each cluster are given above each topography. In addition to the 24 clusters, the mean topography for the outlier cluster is provided.

**Supplementary G.**
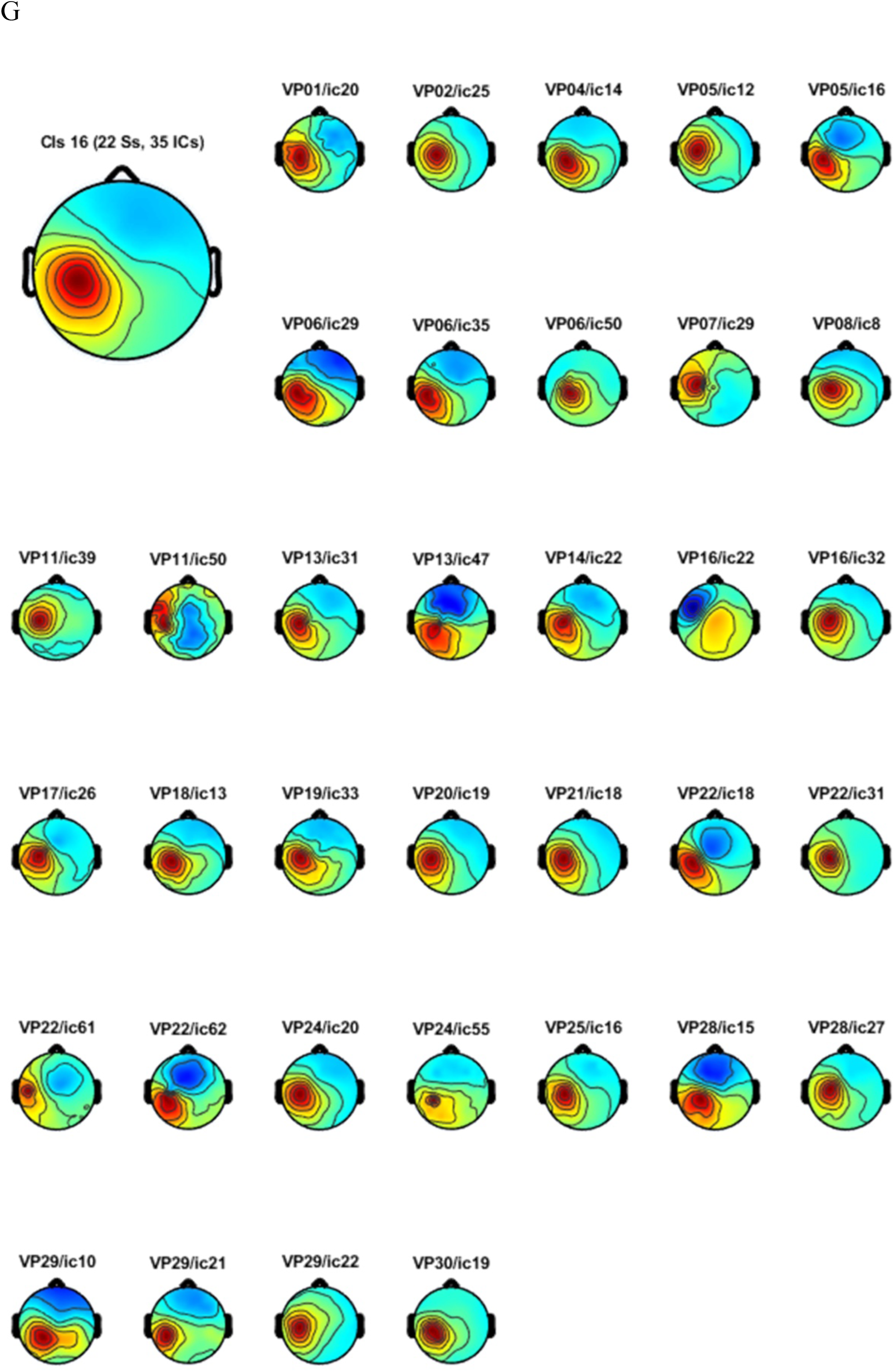
Individual scalp topographies contributing to the contralateral IC cluster in experiment 2.

**Supplementary H.**
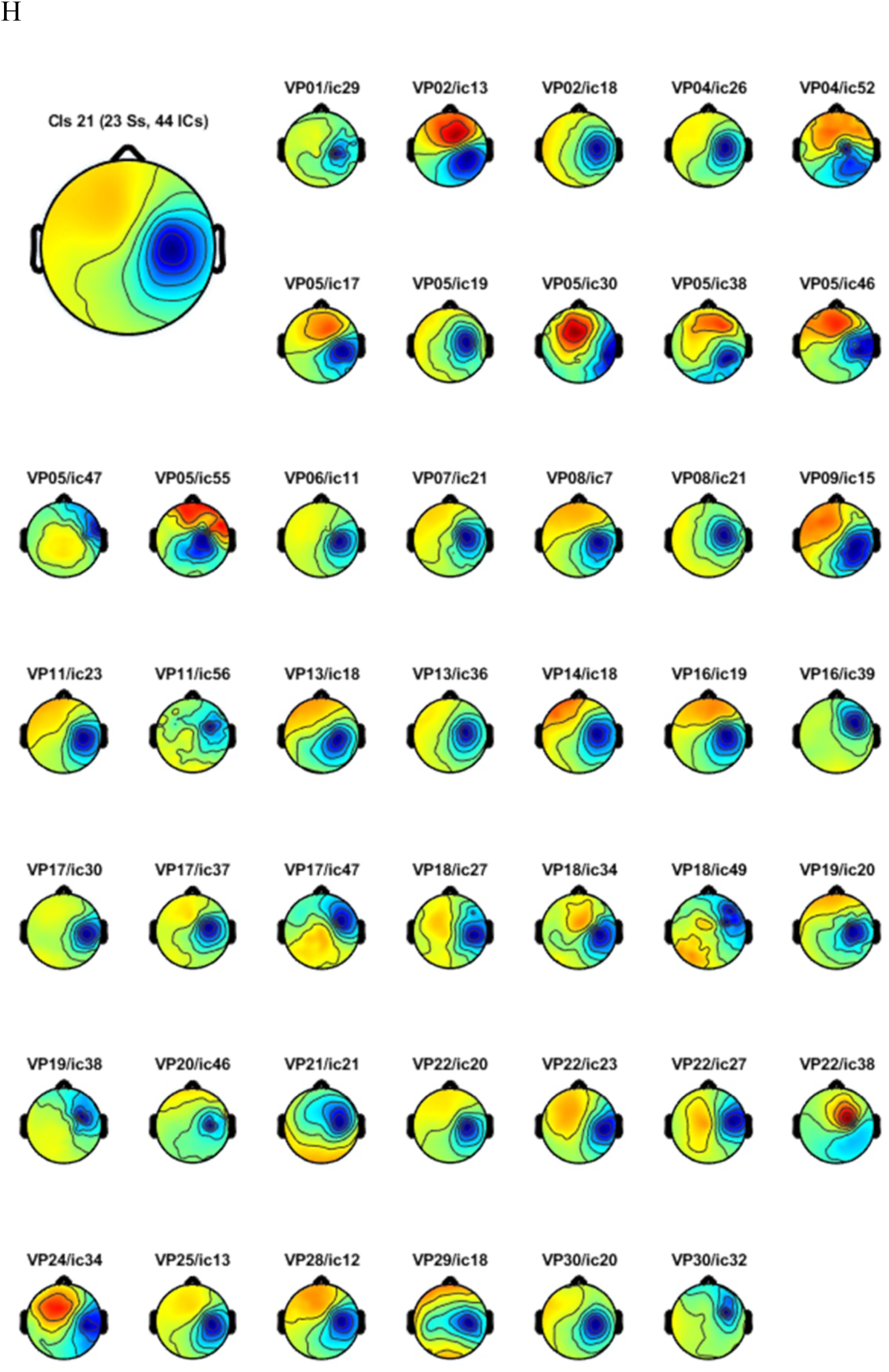
Individual scalp topographies contributing to the ipsilateral IC cluster in experiment 2.

